# Neurogenic archetypes and diversifiers configure the molecular and functional organization of corticofugal output channels

**DOI:** 10.64898/2026.11.24.768951

**Authors:** Shreyas M. Suryanarayana, Yida Zhang, Jessica Zhang, Sweta Srivas, Shengli Zhao, Hemanth Mohan, Yi Li, Z. Josh Huang

## Abstract

Cortical extratelencephalic neurons (ET) derive from either direct neurogenesis from radial glia cells (ET^d^) or indirect neurogenesis through intermediate progenitors (ET^i^), but whether and how neurogenic origin contributes to ET diversity and function remains unknown. Using genetic strategies that differentially target ET^d^ and ET^i^, we profiled their transcriptomes across the isocortex and imaged their activity in motor cortex during a reach-grasp-drink (RGD) task. While both ET^d^ and ET^i^ contribute to all transcriptomically-defined ET subtypes, they diverge significantly in the concerted expression profile of effector-gene families that shape transcriptional regulation, neuronal connectivity and physiology. During the RGD task, motor cortex ET^d^ formed a small yet heterogeneous pool of broadly tuned activity archetypes, while ET^i^ included a diversified set of spatially organized subpopulations that tracked successive actions of RGD. Therefore, neurogenic origin defines a developmental axis of neuronal diversification orthogonal to the current transcriptomic classification scheme, in which ET^i^ differ from ET^d^ transcriptomically through coordinated significant shifts in effector-gene families, while functionally in the motor cortex, they elaborate a small pool of task-generic ET^d^ into specialized output channels that track different phases of goal-directed RGD behavior.

## Introduction

Extratelencephalic (ET) neurons form the principal output channels of the cerebral cortex^1–4^. In contrast to the other two major projection neuron subclass (PN), the intratelencephalic neurons (IT) and corticothalamic neurons (CT), whose axons remain largely within the telencephalon and the thalamus respectively^4^, ET neurons broadcast cortical output to myriad subcortical structures, including the entire subcortical sensory-motor infrastructure^3–5^. Decades of studies have revealed substantial diversity of ET neurons in their projection pattern, physiological property, gene expression, and systems function. For example, single cell transcriptomic profiling has identified multiple clusters within the broad ET class ^6–9^, and single-neuron reconstruction has revealed ET neurons that differ not only in which targets they innervate, but also in their relative innervation strengths to each target^10–12^ . However, the developmental basis and functional significance of ET neuron diversity remain not well understood.

During corticogenesis, ET neurons, like all cortical excitatory projection neurons, are generated either by direct neurogenesis (dNG) from radial glia cells or by indirect neurogenesis (iNG) through *Tbr2*-expressing intermediate progenitors^13–19^. We previously showed that dNG generates all major PN classes while iNG amplifies and diversifies them^13^. We further demonstrated that whereas ET^d^ primarily targets the forebrain and midbrain, ET^i^ dominates the hindbrain (e.g., brainstem and spinal cord), hypothalamic, and neuromodulatory regions with cortical area-specific projections^15^. Furthermore, area-specific spinal projections of ET^i^, but not of ET^d^, emerge from the postnatal pruning of an initial cortex-wide population^15^. These results indicate a substantial expansion and diversification of the ET^d^ projection repertoire by ET^i^. However, the molecular basis and functional significance of ET^d^ versus ET^i^ distinctions remain unclear.

Here, leveraging genetic approaches that differentially target ET^d^ and ET^i^, we profiled their transcriptomes across the isocortex by single-nucleus RNA sequencing and imaged ET neuron activity in the motor cortex using 2-photon microscopy during a forelimb reach-grasp-drink (RGD) task. We show that ET^d^ and ET^i^ both contribute to all transcriptomically-defined ET subtypes yet differ significantly by graded shifts in effector genes; in particular, they diverge in the concerted expression profile of effector-gene families that govern transcriptional regulation, neuronal connectivity and physiology. Furthermore, similar coordinated shifts are also observed in directly and indirectly derived neurons within the CT (CT^d^ and CT^i^) and NP (near-projecting, NP^d^ and NP^i^, respectively) PN subclasses. We thus define an axis of neuronal diversification, orthogonal to current transcriptomic classification of deep-layer PN subclasses in the isocortex. During the RGD task, motor cortex ET^d^ form a small, high-dimensional pool of heterogeneous response-templates that are generic to RGD behavior. ET^i^ reproduce a subset of these ’archetypes’, amplifying several of them one-to-many, and elaborate them into low-dimensional, spatially clustered channels, tuned to distinct successive behavioral phases of RGD.

## Results

### Deep layer PN^d^ and PN^i^ contribute to all transcriptomic subtypes

Given the transcriptomic and anatomic diversity of ET neurons, ET^d^ and ET^i^ may either differentially contribute to different ET transcriptomic subtypes or, alternatively, differentially shape the properties of each subtype. To distinguish these two possibilities, we profiled deep layer PN^d^ and PN^i^ nuclei (deep layer PN expressing *FezF2*) differentially labeled with nuclear membrane-targeted RFP and GFP respectively, across the isocortex by snRNAseq using *FezF2-CreER;Tbr2-Flp;IS-INTACT*^13,15,20^ mice (**Fig. 1a-b; see Methods**). Following quality control, 92,830 FACS-sorted nuclei were retained: 31,550 (34%) from PN^d^ and 61,280 (66%) from PN^i^ libraries. (10 libraries, 17 mice; **Fig. 1c**). Different libraries were consistent in terms of detected genes and transcripts (**Extended Data Fig. 1a**), as well as in subclass compositions (**Extended Data Fig. 1b**). On a joint embedding, PN^d^ and PN^i^ nuclei were intermingled throughout the deep-layer subclasses^21^, L5 ET, L6 CT, L6b and L5 NP (**Fig. 1c**; **Extended Data Fig. 1c-e**). Every transcriptomic type at the finest level consists of both PN^d^ and PN^i^ (**Fig. 1d**), and genes that define deep-layer subtype identity, such as *Bcl11b*, *Tle4*, *Sox5*, *Ldb2* and *Satb2*, were expressed at similar levels in both (**Extended Data Fig. 2a**). We calculated the number of differentially expressed genes between each pair of transcriptomic subtypes at the finest level within PN^d^ versus PN^i^, divided the numbers into nine quantiles based on similarities (that is, higher similarity yields fewer DEG) and plotted their distribution by quantiles (**Fig. 1e**). Both PN^i^ and PN^i^ show large differences in numbers of DEG between clusters; thus, PN^d^ and PN^i^ subtypes span a similar range of transcriptomic differences, indicating comparable diversity and hierarchical organization within the two neurogenic populations.

**Fig. 1.**
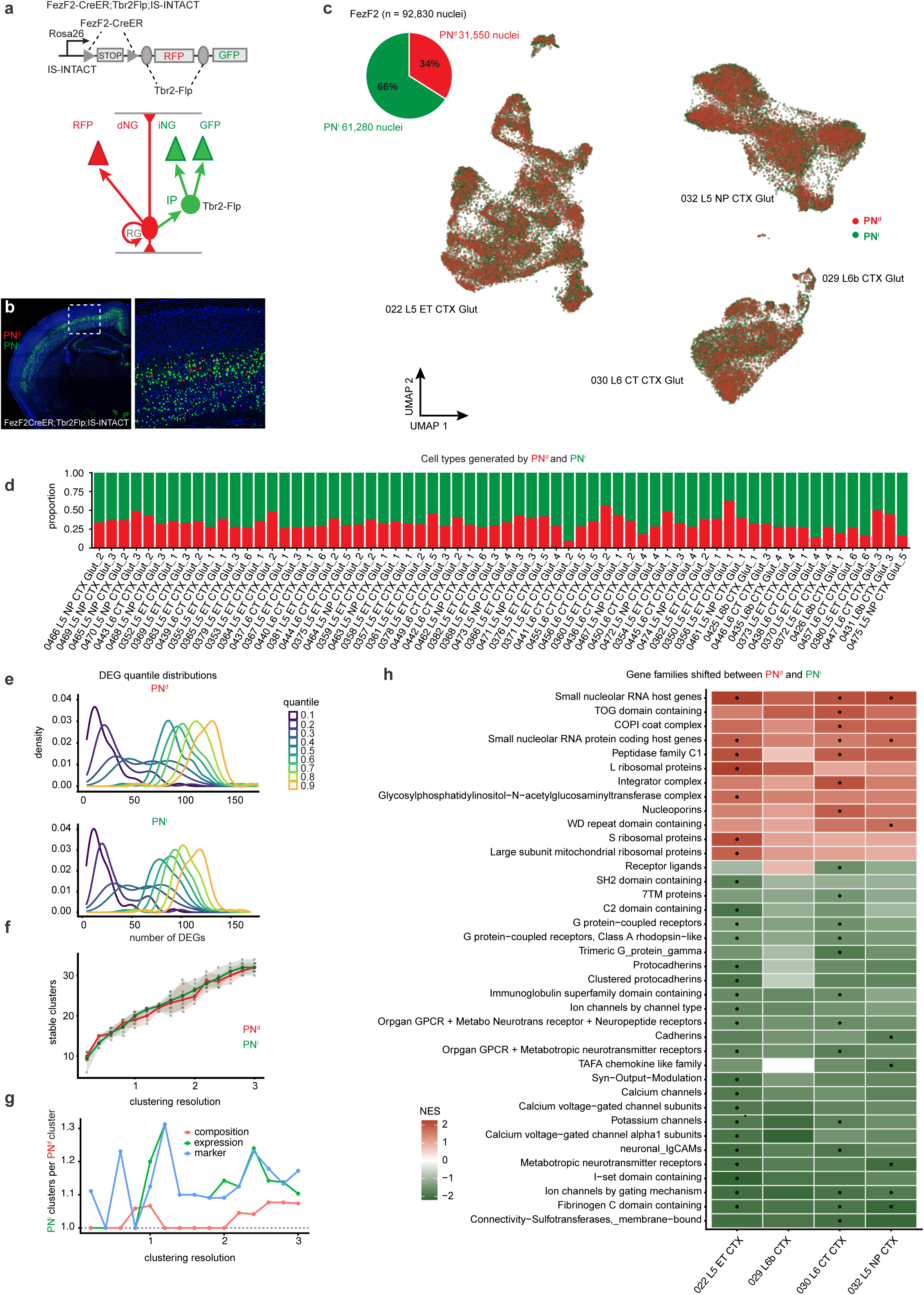
PN^d^ and PN^i^ share a common deep-layer PN transcriptomic identity. **a.** Genetic strategy for labelling dep-layer PN^d^ and PN^i^ nuclei. In *FezF2-CreER*;*Tbr2-Flp*;*IS-INTACT*, *CreER* removes the STOP cassette to express *RFP*, and *Tbr2-Flp* additionally removes *RFP* to express *GFP*, so PN^d^ are *RFP*+ and PN^i^ *GFP*+. **b.** Coronal section from a *FezF2-CreER*;*Tbr2-Flp*;*IS-INTACT* mouse with PN^d^ (red) and PN^i^ (green) nuclei; right, higher magnification. **c.** Composition of *FezF2* nuclei (n = 92,830): PN^d^ 31,550 (34%) and PN^i^ 61,280 (66%); and UMAP of nuclei, PN^d^ (red) and PN^i^ (green) intermingling within the L5 ET, L6 CT, L6b and L5 NP Glut subclasses. **d.** Proportion of each finest-level transcriptomic type (MapMyCells cluster labels as types) generated by ET^d^ (red) and ET^i^ (green); every cluster contains both ET^d^ and ET^i^ in graded, type-specific proportions. **e.** Distribution of the number of DEGs between every pair of clusters within PN^d^ and PN^i^, split into indicated quantiles. The curves show the spread of the number of DEGs between more similar types at 0.1 quantile versus the more distinct types at 0.9 quantile. **f.** Number of stable clusters against clustering resolution for PN^d^ and PN^i^. Lines and colored points, mean across runs; grey points, individual runs; shading, range across runs. Each run clusters a random subsample with equal numbers of nuclei per lineage. **g.** Mean number of PN^i^ groups linked to each PN^d^ group against clustering resolution, with groups linked by cell-type composition, expression or marker overlap; dashed line, one-to-one. **h.** Gene-set enrichment of gene families for PN^d^ versus PN^i^ in each deep-layer subclass. Colour, normalized enrichment score (NES; red, higher in PN^d^; green, higher in PN^i^). Dots, significant gene families in that subclass (FDR < 0.05, same direction in each batch and after sex balancing). Rows, families significant in at least one subclass.

Further, when PN^d^ and PN^i^ nuclei were clustered independently, most clusters in one population had a single counterpart in the other; a few PN^d^ clusters corresponded to two PN^i^ clusters, and only a few clusters in either population lacked a counterpart (**Extended Data Fig. 2b**). The two yielded similar numbers of stable clusters across resolutions (**Fig. 1f**; **Extended Data Fig. 2c**). By expression levels and marker genes^9,21^, PN^i^ also resolved into more than one group per PN^d^ group (up to 1.3) across most resolutions, but not by cell-type composition (at most about 1.07; **Fig. 1g; see Methods**). Thus, both PN^i^ and PN^i^ contribute to all deep-layer transcriptomic subtypes and they span a similar transcriptomic space.

### ET^i^ differ from ET^d^ through coordinated shifts in effector-gene families

Although ET^d^ and ET^i^ contribute to every ET transcriptomic type, differential gene expression analysis revealed robust, substantial, and coordinated differential expression (see **Methods, Supplementary Table 1)** in a set of specific effector gene families. We identified 568 significant differentially expressed genes (DEGs) in L5 ET. Among them, slightly more than half were higher in ET^d^ and the remainder in ET^i^ (**Fig. 2a**). Gene-set enrichment of ET^d^ versus ET^i^ comparison within each subclass (expressed as a normalized enrichment score, NES) gave significant (FDR < 0.05) and reproducible enrichment of gene families (**Fig. 1h, Supplementary Table 2**). In the L5 ET subclass, gene families with higher expression in ET^i^ included connectivity cell-adhesion molecules (neuronal IgCAMs, I-set), ion channels (by gating mechanism, potassium and calcium voltage-gated subunits), metabotropic and modulatory neurotransmitter receptors, G protein-coupled receptors (GPCR) and C2-domain genes; on the other hand, ribosomal, mitochondrial-ribosomal and small nucleolar RNA host-gene families were higher in ET^d^. This gene expression shift however, is at the population level with no single gene family classifying PN^d^ from PN^i^ appreciably above chance (AUROC close to 0.5 in every subclass; **Extended Data Fig. 5a**). Thus, PN^i^ and PN^i^ are distinguished not by categorical markers but by significant, coordinated, population-level shifts across functionally coherent gene families.

**Fig. 2.**
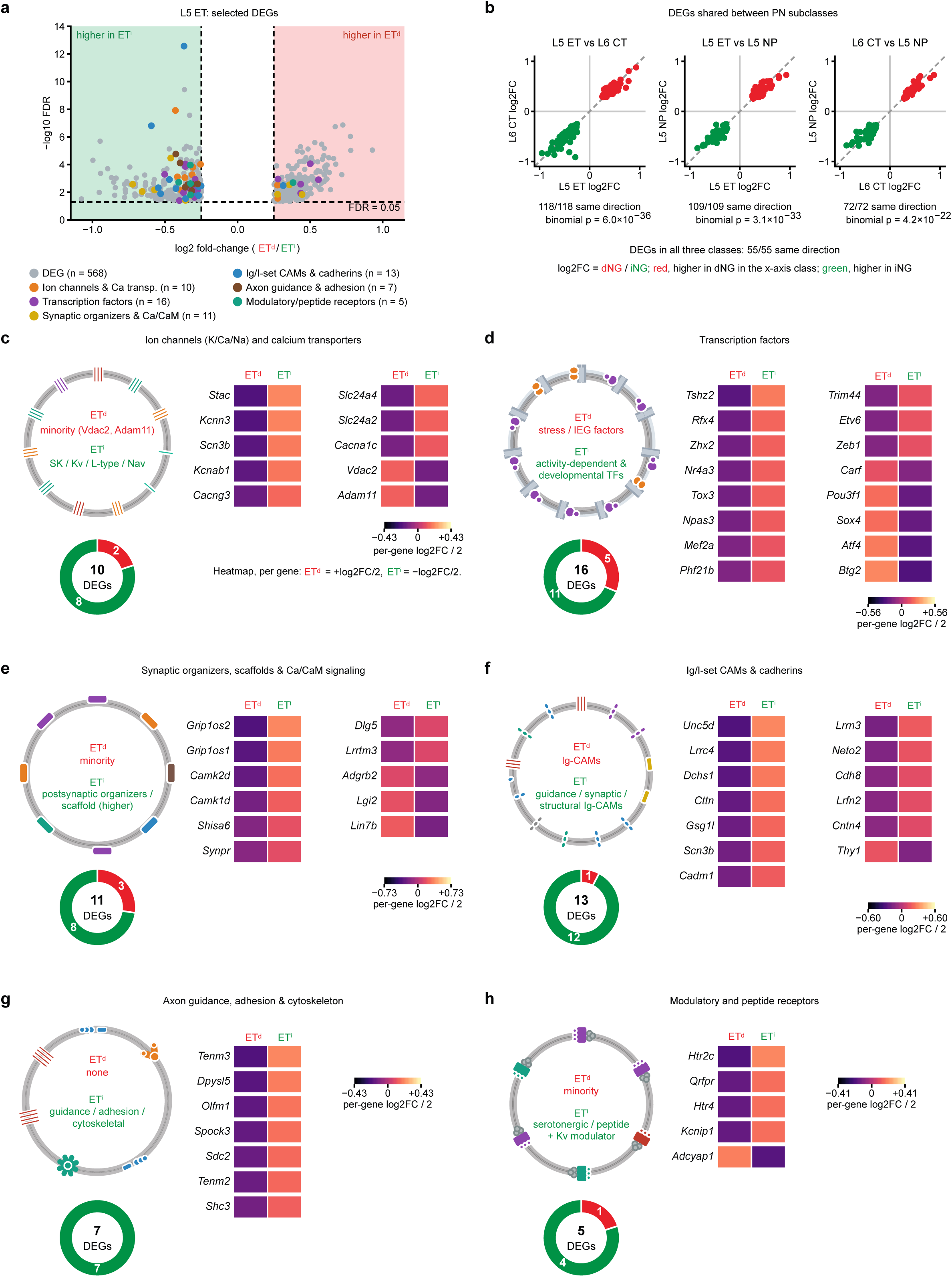
ET^i^ diversifies across effector-gene families. **a.** Volcano of L5 ET DEGs (ET^d^/ET^i^; −log10 FDR versus log2 fold-change). Grey, all 568 DEGs; coloured, DEGs in the selected gene families (ion channels & Ca transporters n = 10, transcription factors n = 16, synaptic organizers & Ca/CaM n = 11, Ig/I-set CAMs & cadherins n = 13, axon guidance & adhesion n = 7, modulatory/peptide receptors n = 5). **b.** DEGs shared between cell classes. log2FC (dNG/iNG) of every gene that is a DEG in both classes of a pair (L5 ET vs L6 CT, n = 118; L5 ET vs L5 NP, n = 109; L6 CT vs L5 NP, n = 72); red, higher in dNG and green, higher in iNG in the x-axis class; dashed line, identity. All shared DEGs change in the same direction (118/118, 109/109 and 72/72; two-sided binomial test against 0.5, p = 6.0 × 10^−3^^6^, 3.1 × 10^−3^^3^ and 4.2 × 10^−2^^2^), as do all 55 DEGs common to the three classes. **c.** Ion channels (K/Ca/Na) and calcium transporters (10 DEGs; 8 higher in ET^i^, 2 in ET^d^). Schematic of the shifted channels on dNG and iNG; donut of the direction split; per-gene heatmap (ET^d^ = +log2FC/2, ET^i^ = −log2FC/2). **d.** Transcription factors (16 DEGs; 11 higher in ET^i^, 5 in ET^d^). ET^i^ carries developmental and activity-dependent factors, ET^d^ the stress/IEG factors; conventions as in c. **e.** Synaptic organizers, scaffolds and Ca²⁺/CaM signaling (11 DEGs; 8 higher in ET^i^, 3 in ET^d^). ET^d^ carries a minority (*Adgrb2*, *Lgi2*, *Lin7b*), ET^i^ the broader repertoire (*Grip1* antisense transcripts *Grip1os1/2*, *Camk2d*, *Camk1d*, *Shisa6*, *Dlg5*, *Lrrtm3*, *Synpr*); conventions as in c. **f.** Ig/I-set cell-adhesion molecules and cadherins (13 DEGs; 12 higher in ET^i^, *Thy1* in ET^d^); conventions as in c. **g.** Axon guidance, adhesion and cytoskeleton (7 DEGs, all higher in ET^i^: *Tenm2*, *Tenm3*, *Shc3*, *Dpysl5*, *Olfm1*, *Spock3*, *Sdc2*); conventions as in c. **h.** Modulatory and peptide receptors (5 DEGs; 4 higher in ET^i^, *Adcyap1* in ET^d^); conventions as in c. For all families, direction is the neurogenic origin with higher expression.

To resolve these family-level differences at the level of individual genes, we examined within the enriched families, gene expression levels consistently shifted toward one neurogenic origin (consistently shifted gene, CSG) in the L5 ET class, and among these, highlighted individual genes with established relevance to cortical development and function. We read these families by the direction and consistency of expression level differences of constituent genes across batches along with individual-gene significance and show those CSGs that are also significant DEGs (**Fig. 2c-h**; see **Methods**). ET^d^ and ET^i^ diverged significantly in transcription factors and effector-gene families for connectivity, ion channels, receptors and intracellular signaling, with most CSGs higher in ET^i^. These shifts were shared across deep-layer PN subclasses: every DEG common to L5 ET and L6 CT (118 of 118), L5 ET and L5 NP (109 of 109) and L6 CT and L5 NP (72 of 72) changed in the same direction (**Fig. 2b**).

Ion channels shifted toward ET^i^ (10 DEGs; **Fig. 2c**), spanning Ca²⁺-activated K⁺ (*Kcnn3*), Kvβ (*Kcnab1*)^22^, L-type Ca²⁺ (*Cacna1c*)^23^, AMPA-auxiliary (*Cacng3*) and Na⁺-channel β (*Scn3b*) subunits, the adaptor *Stac* and the Ca²⁺ transporters *Slc24a2/4*, which shape adaptation, modulation and dendritic Ca²⁺; only the mitochondrial porin *Vdac2*^24^ and *Adam11* were higher in ET^d^. Transcription factors (16 DEGs; **Fig. 2d**) separated by their regulatory role. The eleven higher in ET^i^ were developmental and terminal-selector regulators (*Tshz2*, a deep-layer marker^25^; *Tox3*, *Zeb1*^26^ and *Mef2a*^27^) and activity-dependent factors (*Nr4a3*, *Npas3*^28^), consistent with a broader program for elaborating diverse output identities, whereas the five higher in ET^d^ were stress-responsive and immediate-early regulators (*Atf4*, *Btg2*, *Carf*, *Sox4* and *Pou3f1*^29^), a compact, activity-coupled regulatory core. The strongest effector signal was in Ig/I-set cell-adhesion molecules (Ig-CAMs) and cadherins, which mediate axon guidance, fasciculation and synaptic specificity^30,31^ (**Fig. 2f**). Twelve of thirteen DEGs were higher in ET^i^, including *Unc5d*^32^, *Cadm1*^33,34^, *Cntn4*^35^, *Lrrc4*, *Lrfn2*, *Lrrn3*^36^, *Dchs1*^37,38^ and *Cdh8*^39,40^, and only *Thy1*^41^ in ET^d^. *Unc5d*, a *Tbr2⁺* intermediate-progenitor marker^42^, ranked amongst the most significant DEGs, consistent with a molecular remnant of the indirect neurogenic route. Four of five modulatory and peptide receptor genes were higher in ET^i^ (**Fig. 2h**), including the serotonergic receptors *Htr2c* and *Htr4*^43^, the peptide receptor *Qrfpr* and the excitability-regulating Kv-channel subunit *Kcnip1*^44^, with only *Adcyap1* higher in ET^d^. Of eleven synaptic organizer, scaffold and Ca²⁺/CaM-signaling genes, eight were higher in ET^i^, including *Grip1os1* and *Grip1os2*, antisense transcripts of the *Grip1*^45^ scaffold gene, *Shisa6*^46^, *Lrrtm3*^47^, *Camk2d*^48^ and *Camk1d*, and only *Adgrb2*, *Lgi2*^49^ and *Lin7b* in ET^d^ (**Fig. 2e**). Axon-guidance, adhesion and cytoskeletal genes shifted entirely toward ET^i^ (7 DEGs; **Fig. 2g**): the teneurins^50,51^ *Tenm2* and *Tenm3*, *Dpysl5*^52^, *Shc3*, the proteoglycans *Spock3*^53^ and *Sdc2*^54,55^ and the guidance cue *Olfm1*, a plausible transcriptomic correlate of their expanded, diversified axonal targeting^15^. Thus, ET^i^ differ from ET^d^ mainly in higher expression of genes for wiring, excitability and synaptic signaling. This profile is consistent with their expanded long-range projections^15^ and may shape their distinct output roles across different functional areas of the isocortex.

A similar organization held in the other two deep-layer PN subclasses (**Extended Data Figs. 3-4**). Reading the same six effector families in L6 CT (235 DEGs) and L5 NP (233 DEGs), we again found most family DEGs higher in the iNG-derived population with 46 of 57 in L6 CT and 34 of 48 in L5 NP. Ion channels shifted toward CT^i^ without exception (10 of 10, including *Kcnq5*, *Kcnd3*, *Kcnj6* and *Ryr3*) and predominantly toward NP^i^ (8 of 10, including *Kcnn2*, *Cacna1e* and *Ryr3*), and the same bias held for modulatory and peptide receptors (5 of 5 in L6 CT, including *Htr2c*, *Htr4*, *Grm1* and *Grm7*; 3 of 4 in L5 NP), synaptic organizers and scaffolds (8 of 9 and 6 of 8) and axon-guidance and adhesion genes (8 of 9, including *Tenm3*, *Dpysl5*, *Sema3e* and *Sema5b*; 4 of 6, including *Tenm3* and *Ntng1*) (**Extended Data Figs. 3b,d,f,g** and **4b,d,f,g**). Ig-CAMs and cadherins also leaned toward the iNG population (4 of 7 in L6 CT and 8 of 10 in L5 NP; **Extended Data Figs. 3e, 4e**). Transcription factors were the most balanced family (11 of 17 higher in CT^i^ and 5 of 10 in NP^i^; **Extended Data Figs. 3c** and **4c**), and, as in L5 ET, the stress-responsive factor *Atf4* was higher in the dNG-derived population in both classes. Several genes shifted in the same direction in all three subclasses, including *Grip1os2*, *Camk1d*, *Lrrc4* and *Tenm3* toward iNG and *Atf4* toward the dNG population (see also **Fig. 2b**). Thus, the expanded effector repertoire of iNG-derived neurons is a shared property of deep-layer PN classes.

Furthermore, many of the shifted connectivity, channel, receptor, synaptic and transcription-factor genes are established neurodevelopmental-disorder (NDD) risk genes, we asked whether NDD-risk genes were systematically biased towards one neurogenic origin. Scanning the DEGs of each class and verifying each association (OMIM^56^, SFARI^57^), we identified neuronal NDD-risk genes across epilepsy, autism, intellectual-disability/developmental-delay and psychiatric categories (**Extended Data Fig. 5d-f**). In L5 ET, 18 of 22 NDD-risk DEGs were higher in ET^i^, including the autism genes *Cadm1*^58^ and *Cntn4*^59,60^, the psychiatric risk gene *Cacna1c*^61^, the epilepsy risk gene *Scn3b*^62^, the intellectual-disability risk gene *Lrfn2*^63^, the autism synaptic-adhesion gene *Lrrtm3* and the intellectual-disability kinase *Camk2d*; only the intellectual-disability risk transcription factor *Sox4*^64^, the autism-associated *Gpr85* and *Lin7b* and the neuropeptide gene *Adcyap1* (also a L5 marker) were higher in ET^d^. This ET^i^ bias is significant against the DEG background (82% of NDD-risk DEGs against 46% of the other 546 L5 ET DEGs higher in ET^i^; two-sided Fisher exact test, odds ratio 5.4, p = 8.5 × 10^−4^; **Extended Data Fig. 5d**). The same bias held in L6 CT (24 of 26 NDD-risk DEGs higher in CT^i^, against 52% of the other DEGs; odds ratio 11.0, p = 4.2 × 10^−5^; **Extended Data Fig. 5e**) and in L5 NP (17 of 20 higher in NP^i^, against 44%; odds ratio 7.3, p = 6.3 × 10^−4^; **Extended Data Fig. 5f**), and the 13 NDD-risk genes that were DEGs in more than one class shifted in the same direction in every class (17 of 17 pairwise comparisons; **Extended Data Fig. 5b,c**). Thus, diversification by iNG results in enhanced expression of NDD-risk genes across deep-layer PN classes, providing evidence for a plausible link of neurogenic origin to the genetic architecture of neurodevelopmental disease.

### ET^d^ forms sparse, high-dimensional activity templates

The consistent and coordinated transcriptomic differences between ET^d^ and ET^i^, with ET^i^ carrying an expanded repertoire of connectivity, channel, receptor and synaptic effectors, may differentially craft the anatomical and physiological properties of these subpopulations, to shape and diversify their function in behavior. To test this possibility, we recorded ET^d^ and ET^i^ activity in behaving mice. We used an intersectional strategy that converts a transient developmental signal into a permanent, virally targetable handle which allows viral-access to ET^d^ and ET^i^^15^. In *FezF2TC;Tbr2Flp* double-allele mice, ET^d^ constitutively express *tTA*, whereas those passing through a *Tbr2*^+^ intermediate progenitor (ET^i^) express CreER^15^. Spectrally separable indicators can therefore be assigned to each subpopulation in the same animal by co-injecting *AAV-TRE-jRGECO1a*^65^ (ET^d^) and *AAV-DIO-jGCaMP8f*^66^ (ET^i^) into secondary motor cortex^15^ (a major active hub in RGD^67^; **Fig. 3a**). ET^d^ and ET^i^ were imaged simultaneously in layer 5b while the mouse performed a head-fixed RGD task, with two synchronized cameras recording the forelimb and face (**Fig. 3b**; **Extended Data Fig. 6d**; see also^67^). In each session, the populations were intermingled, and after segregating the two populations, ET^d^ was smaller with 197 (9.8%) against 1,816 ET^i^ (90.2%; **Fig. 3c,d**; **Extended Data Fig. 6a**; n = 4 mice; 17 sessions; see also^15^). A separate cohort labelled the entire ET subclass (*pan-FezF2*, using *FezF2CreER*^20^) with a single indicator (*GCaMP8m*^66^), giving a pan-ET reference (734 ET^all^, n = 3 mice; 10 sessions).

**Fig. 3.**
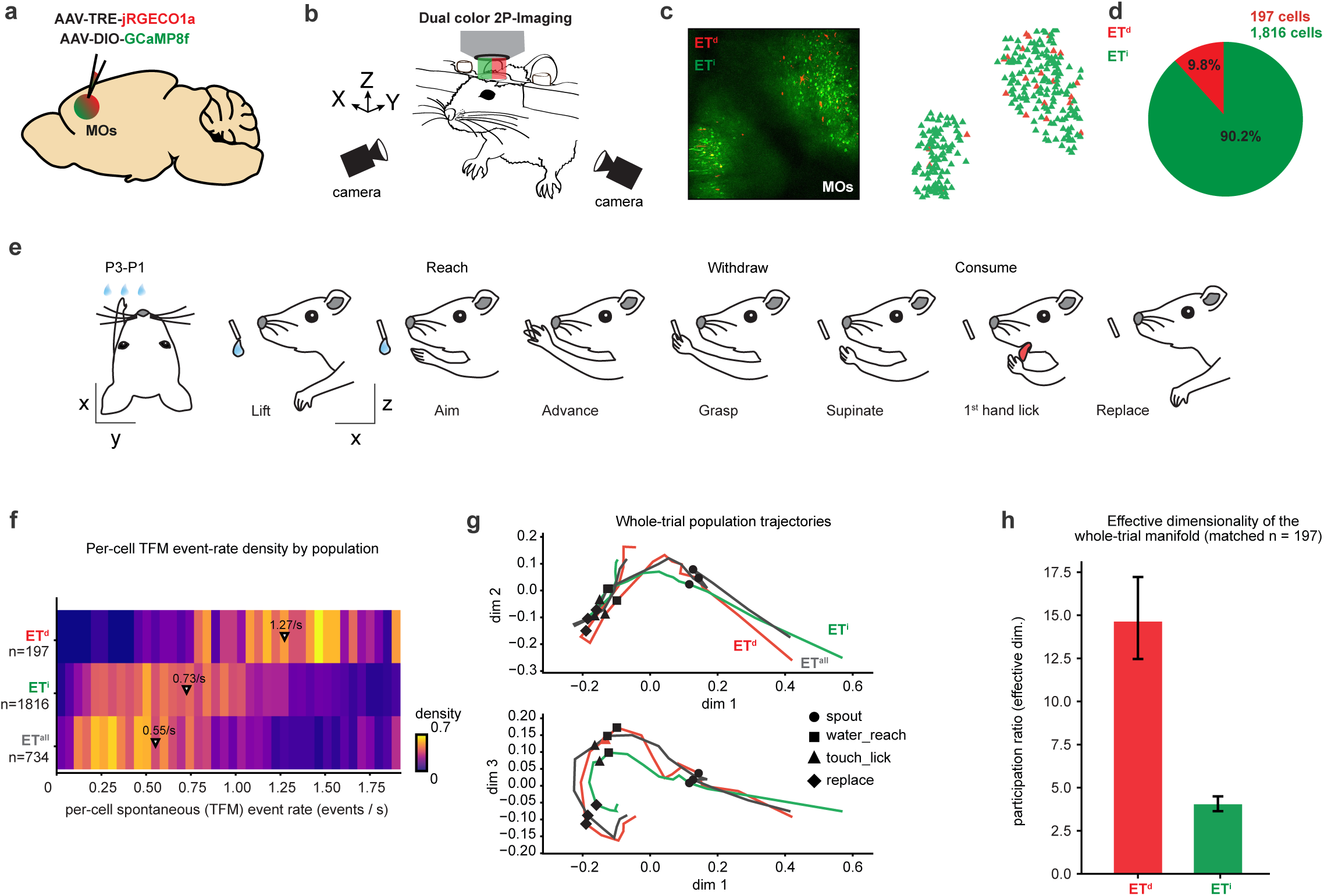
Simultaneous two-color imaging of ET^d^ and ET^i^ reveals a sparse, high-dimensional and heterogeneous ET^d^ population. **a.** Co-injection of *AAV-TRE-jRGECO1a* (ET^d^) and *AAV-DIO-GCaMP8f* (ET^i^) into MOs of *Fezf2TC;Tbr2Flp* mice, in which ET^d^ express tTA while ET^i^ excise tTA and express CreER, allowing independent viral access to the two populations. **b.** Dual-color two-photon imaging of ET^d^ and ET^i^ with two synchronized forelimb and face cameras during head-fixed RGD behavior. **c.** Left, example field of view (fov) in MOs with intermingled ET^d^ (red) and ET^i^ (green) somata; right, soma map of the same field. **d.** Composition of the imaged population: 197 ET^d^ (9.8%) and 1,816 ET^i^ (90.2%) cells. **e.** Trial structure of RGD. The spout advances to one of three positions (P1–P3), followed by jittered water delivery, lift, aim, advance, grasp, supinate, first hand lick and replace. **f.** Density heatmap of per-cell spontaneous (TFM) event rates for ET^d^ (n = 197), ET^i^ (n = 1,816) and ET^all^ (n = 734); medians marked (1.27, 0.73 and 0.55 events/s). **g.** Whole-trial population trajectories for ET^d^, ET^i^ and ET^all^, Procrustes-aligned into a common frame and shown as dim 1 against dim 2 (top) and dim 1 against dim 3 (bottom). Symbols mark four behavioral events along the trial (spout, water/reach, touch/lick and replace). **h.** Effective dimensionality (participation ratio) of the whole-trial response manifold for ET^d^ and ET^i^ at matched cell count (n = 197). Bars, participation ratio; error bars, 95% bootstrap intervals over matched-n draws (14.6 [12.5–17.2] versus 4.0 [3.6–4.5]; ET^d^ exceeded ET^i^ in 100% of matched-n draws).

During the task, the spout advances to one of three positions (**Extended Data Fig. 6c**), water is delivered after a variable delay, and the mouse lifts its hand, aims, advances, grasps, supinates, licks the water from its hand and replaces the paw (**Fig. 3e**; **Supplementary Video 1**, also see^67^). We first examined spontaneous activity of ET^d^ and ET^i^ during task-free windows when mice were not engaged in RGD actions (task-free movement,TFM; **Fig. 3f; Extended Data Fig. 6f**; see **Methods**). ET^d^ showed a higher median spontaneous transient rate than both ET^i^ and ET^all^ (1.27, 0.73 and 0.55 events/s respectively). Populations were subsampled to a matched n = 197 (number of ET^d^) over 50 draws and compared by permutation; all three pairwise differences reached the resolution limit of that test (Bonferroni p = 0.015 in each case; **Extended Data Fig. 6g**), and ET^d^ rates were broadly distributed across the ET^i^ range. The same held during the RGD window, where ET^d^ were again more active than ET^i^ and ET^all^ (1.17, 0.68 and 0.55 events/s; Bonferroni p = 0.015 for all three comparisons; **Extended Data Fig. 6h**). ET^d^ thus is a small and more active population whose activity patterns span the breadth of the larger ET^i^.

To ask whether the two populations traverse different activity states, we traced whole-trial population trajectories through state space (**Fig. 3f**) and computed disparities at a matched cell count. ET^d^ and ET^i^ differed by 0.135 (IQR 0.125 -0.143) against a floor of 0.026 and a null of 0.009 (permutation p = 0.005), clearing both references. The ET^all^ population was likewise distinct from each (ET^d^ - ET^all^ 0.073, floor 0.033; ET^i^ - ET^all^ 0.123, floor 0.020; both p = 0.005) and lay between them, as expected of a population that contains a mix of both (**Extended Data Fig. 6e**). Thus, ET^d^ and ET^i^ follow different sequences of activity states across TFM and RGD, with ET^d^ being a sparse and functionally broad population. The two populations also occupied response manifolds of markedly different effective dimensionality. At matched cell count, the whole-trial participation ratio was roughly 3.6-fold higher for ET^d^ than ET^i^ (14.6 [95% CI 12.5 - 17.2] versus 4.0 [3.6 - 4.5]; non-overlapping bootstrap intervals; ET^d^ exceeded ETⁱ in 100% of matched-n draws; **Fig. 3g**), computed on the same whole-trial activity profiles as the trajectory analysis. Thus, ET^d^ occupies a high-dimensional response manifold whereas ET^i^ is comparatively low-dimensional. The difference held within the single-indicator ET^all^ cohort, where cells matched individually to ET^d^ had a higher participation ratio than cells matched to ET^i^ at matched cell count (n = 40; cell-to-cell matching, see below; **Extended Data Fig. 6i**).

A central tenet to examine is, if ET^d^ are high-dimensional with heterogenous activity profiles, do the larger ET^i^ simply amplify them or are there differences? We asked how the response repertoires of the two populations relate to each other. For each neuron we formed a response profile, the trial-averaged dF/F_0_ across the full trial (**Fig. 4a**). An ordered cell-cell correlation shows that ET^i^ varies as a single smooth gradient while ET^d^ is more heterogeneous, and neither resolves into discrete blocks (**Fig. 4b**). We therefore related the two populations cell by cell. Each ET^i^ cell was matched to the ET^d^ cell with the most similar profile over the RGD window (−2 to +10 s from spout onset; Pearson r), counting a match when r exceeded 0.79, the 95th percentile of a circular time-shift null. By this criterion, 285 of 1,816 ET^i^ cells (16%) reproduced the profile of an ET^d^ cell (median r = 0.86), and they converged on only 18 of the 197 ET^d^ cells (**Extended Data Fig. 7a**). The mapping was one-to-many, with each ET^d^ cell converging on average to 15.8 ET^i^ cells (median 4), 14 of the 18 received more than one, and the largest fan, 104 ET^i^ cells, converged on an ET^d^ cell whose activity rose slowly through RGD. The remaining 1,531 ET^i^ cells (84%) had no ET^d^ counterpart, and 179 of the 197 ET^d^ cells (91%) were reproduced by no ET^i^ cell. ET^i^ therefore amplifies a few ET^d^ response profiles, several of them one-to-many, while generating many profiles that ET^d^ lacks. The same structure held in the single-indicator ET^all^ cohort. Matching each ET^all^ cell to its most similar ET^d^ or ET^i^ cell assigned 56 of 734 (7.6%) to ET^d^, close to the 9.8% measured by neurogenic labelling (binomial p = 0.054), and these ET^d^-matched cells again spanned a larger region of response-profile space than ET^i^ -matched cells (reference-shuffle p = 0.005; **Extended Data Fig. 7d,e**), showing that the difference is not simply contingent on indicator difference. Further, whole-population dispersion, the mean distance between the response profiles of two cells of the same neurogenic population, was greater in ET^d^ than in ET^i^ (29.9 versus 22.7) in 100% of matched-n bootstrap draws (median difference 7.26, 95% CI 6.43-8.00; p < 0.001; **Extended Data Fig. 7b,c**). Taken together, ET^d^ is a sparse, high-dimensional and heterogeneous population, while ET^i^ builds on ET^d^, amplifying some of its response profiles while also generating novel ones. Further, dispersion estimated from random subsets of ET^d^ reached its full-population value within about 20 cells, and ET^d^ exceeded ET^i^ at every matched subset size down to 10 cells (100% of draws), so the 197 ET^d^ neurons are a near-complete sample of a genuinely small heterogeneous population (**Extended Data Fig. 7f,g**).

**Fig. 4.**
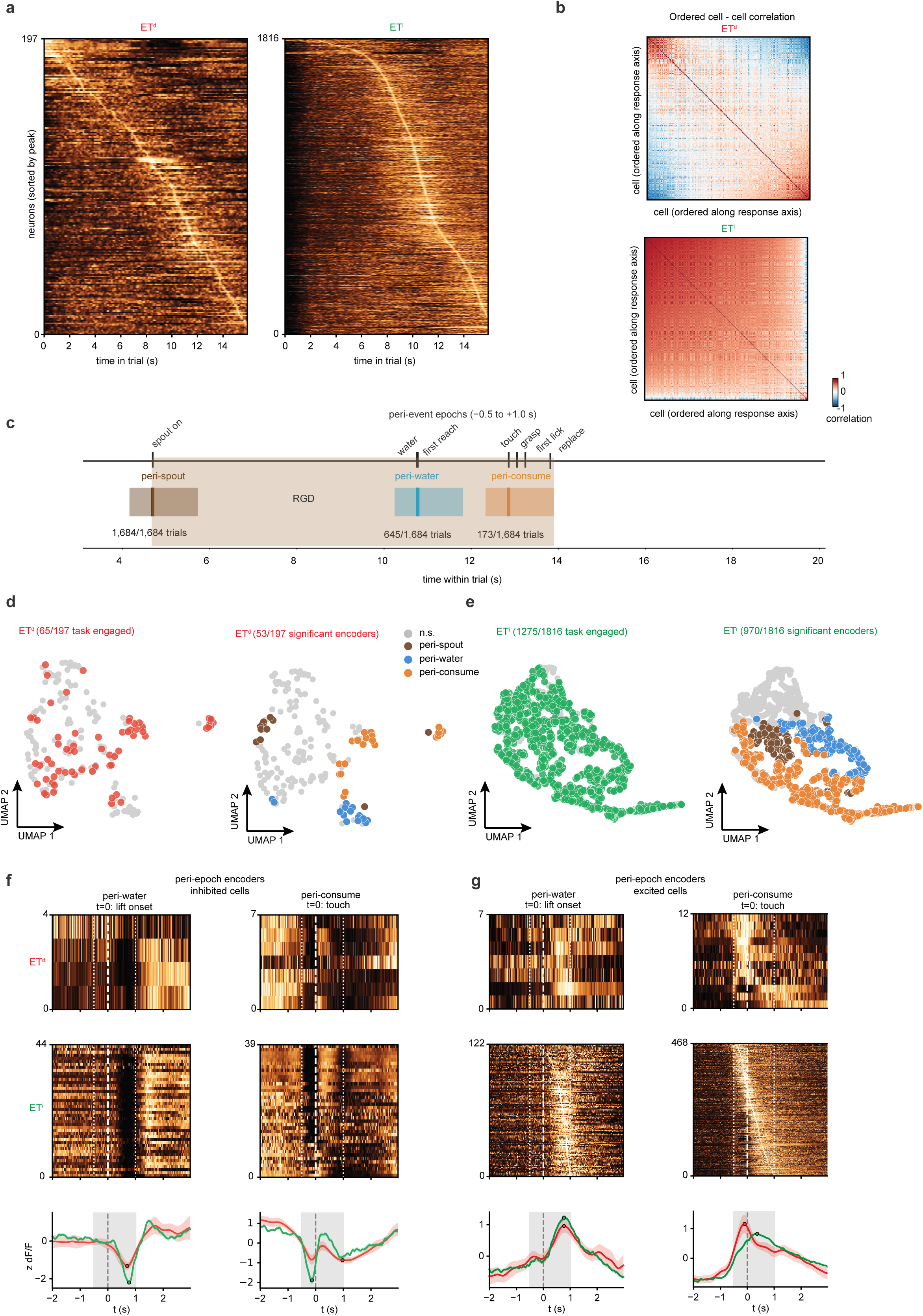
ET^d^ forms heterogeneous activity types while ET^i^ resolves into behavior-specific output channels. **a.** Trial-averaged whole-trial activity of every ET^d^ (n = 197) and ET^i^ (n = 1,816) cell, sorted by time of peak activity. **b.** Ordered cell–cell correlation of whole-trial response profiles for ET^d^ and ET^i^, cells ordered along the dominant response axis; colorbar: Pearson r. ET^i^ varies as a single smooth gradient while ET^d^ is more heterogeneous; neither resolves into discrete blocks. **c.** Peri-event epochs used for the single-cell analysis in **d-g** (peri-spout, peri-water and peri-consume), each −0.5 to +1.0 s about its anchoring event (spout on, first reach after water delivery and first touch after water delivery), with usable-trial counts; shading, RGD window. The four epochs used for single-cell decoding, including peri-replace, are shown in **Extended Data Fig. 8a**. **d.** UMAP embedding of single-cell peri-epoch encoding for ET^d^. Left, task-engaged cells (RGD versus TFM, AUROC > 0.5, p < 0.05; 65/197) against remaining cells (grey); right, significant peri-epoch encoders (53/197) colored by preferred epoch; grey, n.s. **e.** As in **d** for ET^i^ (task-engaged, 1,275/1,816; peri-epoch encoders, 970/1,816). **f.** Peri-epoch encoders whose activity decreased about the epoch anchor (inhibited), for peri-water (t = 0, first reach after water delivery, detected as lift onset) and peri-consume (t = 0, touch). Trial-averaged activity of every ET^d^ (top; 4 and 7 cells) and ET^i^ (middle; 44 and 39 cells) encoder, each row z-scored and sorted by time of trough; bottom, mean ± s.e.m. (ET^d^ red, ET^i^ green). dotted lines, epoch window (−0.5 to +1.0 s); dashed line, anchor; circles, trough. Peri-spout encoders are not shown. **g.** As in **f** for encoders whose activity increased (excited; ET^d^ 7 and 12 cells, ET^i^ 122 and 468 cells); rows sorted by time of peak; circles, peak.

### ET^i^ creates behavior phase-specific output channels

If ET^i^ elaborates the ET^d^ activity repertoire, its response types could possibly distinguish stages of the RGD action that ET^d^ do not. We divided the RGD task into four separable peri-event epochs, peri-spout, peri-water, peri-consume and peri-replace, spanning 0.5 s before to 1.0 s after their respective anchoring events (spout movement, first reach post water delivery, first touch post water delivery, replace onset respectively; **Fig. 4c**; **Extended Data Fig. 8a**); because RGD is fast and behavioral events can crowd together (see **Extended Data Fig. 6b**), an epoch contributes to a trial only when its window does not overlap a neighbor, so the peri-spout epoch is usable on every trial, peri-water on 645, peri-replace on 463 and peri-consume on 173 of 1,684 trials (**Extended Data Fig. 8a**). For each cell we took its mean activity in each peri-epoch on each trial, learned the four epoch means from all but one trial, and asked which epoch the left-out trial’s values matched best, with accuracy referenced to a within-trial label shuffle. Significantly more ET^i^ than ET^d^ cells decoded the epochs above the shuffle null, 721 of 1,816 (39.7%) against 38 of 197 (19.3%; Fisher exact p = 6.8 × 10^−9^; **Extended Data Fig. 8b**), both well above the 5% expected by chance. Deconvolution, to control for a possible indicator confound in dual dF/F_0_, left the difference intact (528 of 1,816, 29.1%, against 24 of 197, 12.2%; p = 8.6 × 10^−8^; **Extended Data Fig. 8b**). Epoch-decoding cells of both ET^d^ and ET^i^ covered all four peri-epochs, and on dF/F_0_ their preferred-epoch split did not differ (χ^2^ p = 0.71; **Extended Data Fig. 8c**).

A single-cell encoding analysis gave the same contrast (**Fig. 4d,e**). Here, we used the three peri-epochs leading up to consumption (**Fig. 4c**), because the end of the action, the peri-replace, is analyzed separately (see below; **Extended Data Fig. 12**). Most ET^i^ cells were engaged by the task, being more active during RGD than during TFM (1,275 of 1,816, 70%), and over half encoded a specific peri-epoch (970, 53%), with cells preferring the same peri-epoch grouping together (**Fig. 4e**). In ET^d^, both kinds of cell were less common (65 of 197, 33%, engaged; 53, 27%, encoding a peri-epoch) and were scattered (**Fig. 4d**). In both, encoders either decreased or increased their activity about the anchor of their preferred epoch (**Fig. 4f,g**). ET^i^ is thus both recruited by the task and divides it into peri-epochs, whereas ET^d^ represents RGD more generally. Peri-epoch information was also widespread in the single-indicator pan-FezF2 ET^all^ population,and on spike rates inferred with a GCaMP8m-specific CASCADE model^68^, 249 of 734 ET^all^ cells (33.9%) decoded the four peri-epochs above null (**Extended Data Fig. 8b**). Thus, ET^d^ form a generalist population in which few cells are devoted to any one peri-epoch, whereas ET^i^ diversify into peri-epoch-specific output channels that encode distinct RGD phases.

### A basal-ganglia action-selection model cleanly sequences RGD from ET^i^ activity but not ET^d^

We next asked whether ET^i^ peri-epoch activity is organized well enough to drive the ordered RGD action sequence. Because targeting specific ET^i^ peri-epoch channels for optogenetic manipulation is not feasible, we instead asked whether ET^d^ and ET^i^ differ in how well an established model of basal-ganglia action selection^69,70^ can use their activity to sequentially orchestrate RGD. In the model, a focused striatal direct pathway (D1 spiny projection neurons) competes against a diffuse subthalamic off-surround scaling with total salience across channels, and the output nuclei (SNr/GPi) tonically inhibits its targets and releases the winner by disinhibition (see also^69^; **Extended Data Fig. 9a**; **Methods**). The model was given three channels, one per behavioral peri-epoch (peri-water, peri-consume and peri-replace; peri-spout was excluded because spout arrival is not a motor phase), with each channel’s salience taken from the deconvolved activity of the whole recorded population, each cell weighted by its mean response in that peri-epoch (**Extended Data Fig. 9b**), and the readout asked whether the channel selected at each moment of RGD progression, was the one whose peri-epoch was current (chance 1/3). Both ET^d^ and ET^i^ sequenced RGD well above chance, the correct channel selected during 95.1% of the driven timeline with ET^i^ against 79.3% with ET^d^ (**Extended Data Fig. 9c**), each channel winning in turn across the event-anchored trial; however, ET^i^ selected the correct peri-epoch channel sequentially while ET^d^ showed noisy and multiple channel selection (**Extended Data Fig. 9d,e**). Thus, ET^i^ supplies independent peri-epoch channels that a basal-ganglia selector can sequence into ordered behavior.

### ET^i^ output channels are spatially organized in the cortical sheet

Since ET^i^ forms peri-epoch-specific output channels, we asked whether cells sharing an epoch preference are spatially organized in the cortical field of view (fov). Each ET^i^ cell that decoded the four peri-epochs above its shuffle null (721 of 1,816; **Extended Data Fig. 8b**) was labelled by its preferred peri-epoch, and we asked whether its nearest neighbors shared that preference more often than expected. Each field was tested against a shuffle of its own labels, since each is an independent fov (**Fig. 5a**). Of 17 fields (imaging sessions), 10 carried enough cells to test (at least 25); 8 of these 10 exceeded their own null at 12 nearest neighbors and 3 reached significance individually (Fisher combined p = 2.1 × 10^−3^; **Fig. 5c**). Coloring each cell by the peri-epoch dominating its immediate neighborhood reveals local patches sharing a preference, recurring across fields (**Fig. 5a**). The arrangement is local, as concordance exceeded the shuffle null in most fields at every neighborhood size tested (7, 6 and 8 of 10 fields at 1, 5, 8 and 12 nearest neighbors), and the combined test became significant as more neighbors were averaged (p = 0.12, 0.067, 0.014 and 0.002; **Fig. 5b**), the 8th and 12th neighbors lying at a median of 113 and 148 µm. ET^i^ cells preferring the same peri-epoch therefore sit near one another, so the epoch-specific channels are locally organized in MOs and recur across different fov’s. ET^d^ appear salt-and-pepper on the same maps, with only 38 epoch-decoding ET^d^ cells across all fields, no field reaches the number of cells needed to estimate concordance (**Fig. 5a**).

**Fig. 5.**
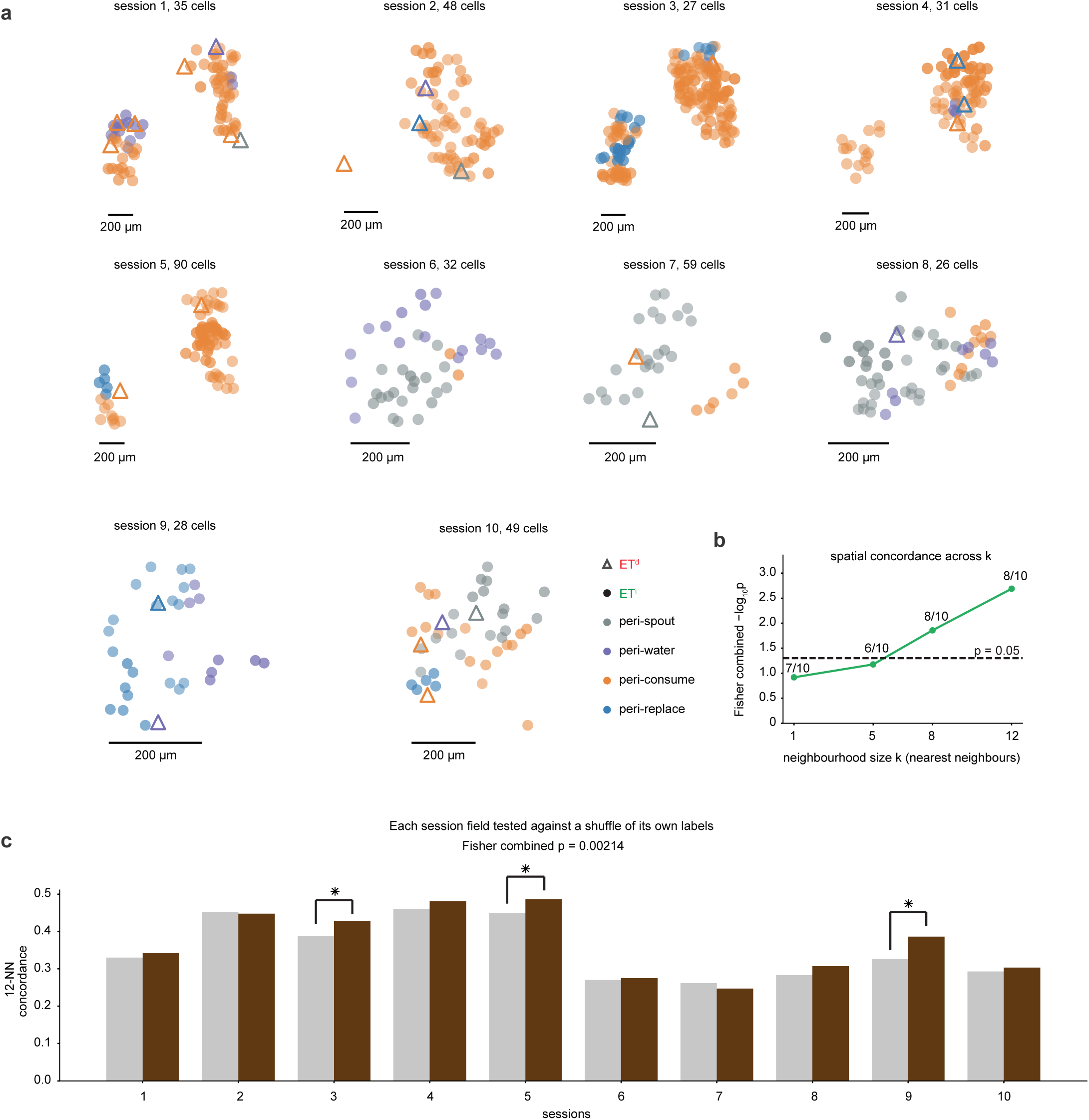
ET^i^ output channels are spatially organized in the cortical sheet. **a.** Per-session soma maps of ET^i^ cells that decode the four peri-epochs above their shuffle null (dF/F_0_; **Extended Data Fig. 8b**), for the 10 fields of view with at least 25 such cells. Each cell is colored by the peri-epoch dominating its neighborhood (its 12 nearest cells and itself), with opacity scaled by that epoch’s local excess over its field-wide frequency; triangles, above-null ET^d^ cells colored by their own preferred epoch. Each panel is an independent field of view (session); scale bars, 200 µm. **b.** Spatial concordance across neighborhood size k (1, 5, 8 and 12 nearest neighbors): Fisher-combined −log_10_ p across the 10 fields; numbers, fields above their own null; dashed line, p = 0.05. **c.** 12-nearest-neighbor concordance per field, observed (brown) against a within-field label shuffle (grey); 8/10 fields above null, 3 individually significant (asterisks, p < 0.05; Fisher combined p = 2.1 × 10^−3^).

### ET^d^ and ET^i^ both encode RGD task kinematics, while ET^i^ extends it to spontaneous movements

Having shown that ET^i^ diversifies ET^d^ into spatially-clustered peri-epoch channels, we asked whether the two populations nonetheless share a common function, kinematic coding, a hallmark of cortical motor areas^71–73^ and ET neurons^6,67^. We fitted a cross-validated encoding model to every cell using 18 continuous kinematic series spanning hand position, speed and path length, aiming, digit aperture, hand-to-spout and hand-to-mouth distance, and orofacial and water-related variables (**Extended Data Fig. 10**; see **Methods**). Both ET^d^ and ET^i^ encoded task kinematics strongly. In the RGD window, 40.1% of ET^d^ (79 of 197) and 49.1% of ET^i^ (891 of 1,816) cells met the encoding criterion, far above the 5% expected by chance, with ET^i^ the higher (Fisher exact p = 0.02; **Fig. 6a**). Strong encoding was also seen in ET^all^ (65.4% of cells; **Extended Data Fig. 11a,e**) and encoding cleared chance in every ET^i^ session and in most ET^d^ sessions (**Extended Data Fig. 11g**).

**Fig. 6.**
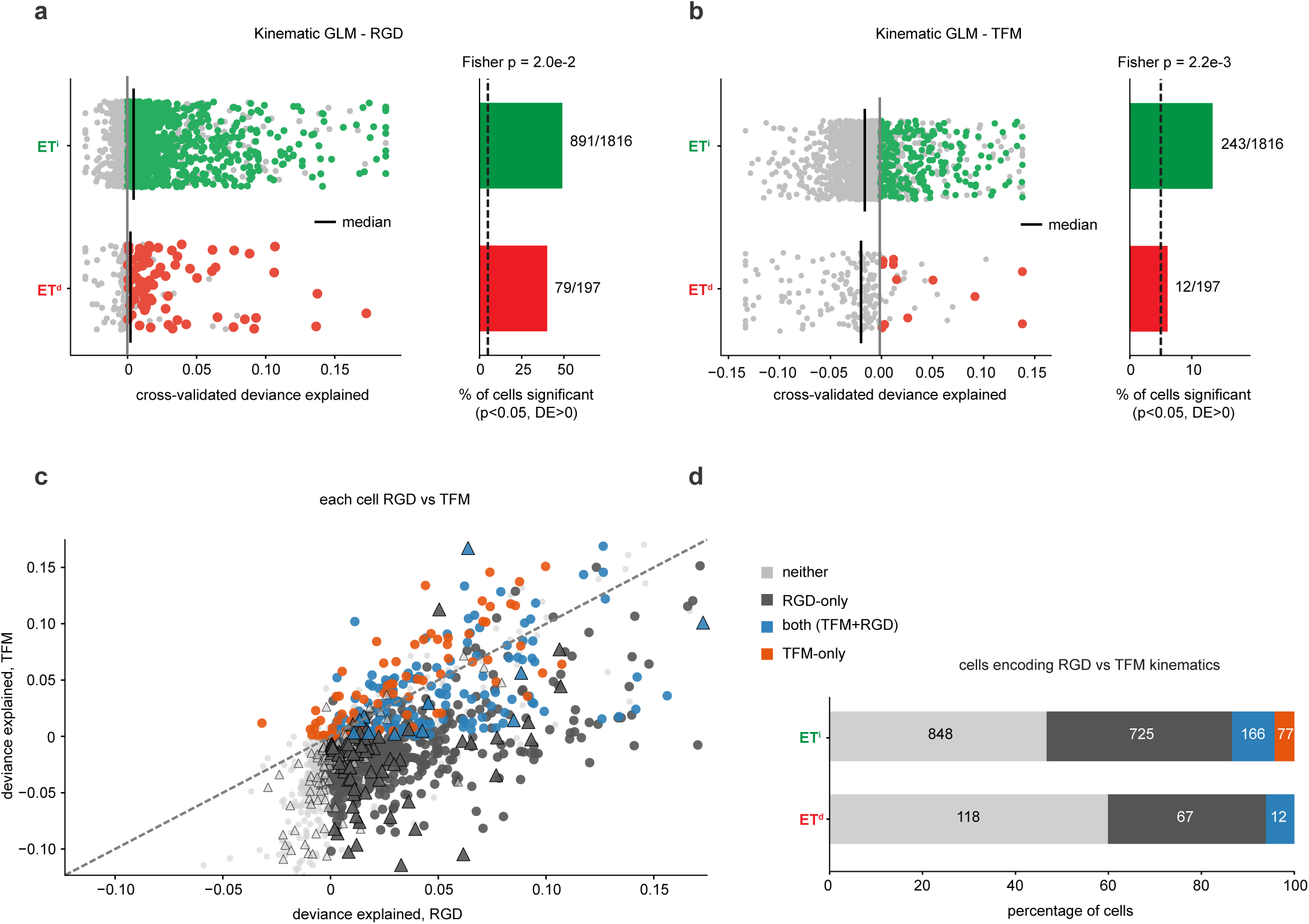
Both ET^d^ and ET^i^ encode RGD task kinematics, and ET^i^ extends this coding to spontaneous movement. **a.** Per-cell kinematic GLM (18 series, five lags), RGD. Left, cross-validated deviance explained per cell for ET^i^ (n = 1,816) and ET^d^ (n = 197); colored, cells encoding above the circular-shift null (p < 0.05, DE > 0); grey, others; vertical lines, medians. Right, percentage of cells encoding (ET^i^ 891/1,816, 49.1%; ET^d^ 79/197, 40.1%; Fisher exact p = 0.02); dashed line, chance (5%). **b.** As in **a** for TFM (ET^i^ 243/1,816, 13.4%; ET^d^ 12/197, 6.1%, at chance, binomial p = 0.28; Fisher exact p = 2.2 × 10^−3^). **c.** Deviance explained per cell in RGD against TFM; circles, ET^i^; triangles, ET^d^; colored by encoding category as in **d**; dashed line, identity. **d.** Encoding category per cell: neither window, RGD only, both (TFM + RGD) or TFM only (ET^i^ 848/725/166/77; ET^d^ 118/67/12/0).

However, ET^d^ and ET^i^ differed during TFM. Refitting the same model on the same cells during TFM (periods of spontaneous movement unrelated to RGD), collapsed encoding in ET^d^ while leaving it in place in ET^i^, with the proportion of encoding cells decreasing from 40.1% to 6.1% in ET^d^ and from 49.1% to 13.4% in ET^i^ (Fisher exact p = 2.2 × 10^−3^; **Fig. 6b**), a reduction also seen in ET^all^ (65.4% to 11.3%; **Extended Data Fig. 11a,b,f**). At 6.1% (12 of 197), ET^d^ is indistinguishable from chance (binomial p = 0.28) and its kinematic encoding disappears into the null, whereas ET^i^ does not collapse, since 243 of 1,816 cells continue to encode, nearly three times chance (**Fig. 6b**). Cell by cell, ET^i^ spans task-bound cells that encode RGD but not TFM (725), cells that encode both (166) and those that encode TFM alone (77; **Fig. 6c,d**). In ET^d^, no cell encodes TFM without also encoding RGD (0 of 197), whereas in ET^i^ about a third of the 243 TFM-encoding cells do so, an additional class for which ET^d^ has no counterpart (**Fig. 6d**). In ET^all^, most encoding cells encoded RGD alone (401) or both windows (79), and 4 encoded TFM alone (**Extended Data Fig. 11c,d**). Kinematic coding in ET^d^ is therefore bounded by the task, whereas ET^i^ diversifies it beyond the task in two ways, with most TFM encoders also encoding RGD, carrying their coding into spontaneous movement, while another distinct subset encoding TFM alone.

### ET^i^ forms separate output channels for a proactive anticipatory and a reactive terminal ending of the RGD action

The RGD behavior can end in one of two ways. On most trials, the mouse returns the hand to the perch immediately after consumption, a reactive terminal movement that closes the sequence (**Supplementary Video 2**); on some trials, it keeps the hand raised instead, a proactive posture, ’hold’, in anticipation of more water (**Extended Data Fig. 12a**; **Supplementary Video 3**). Both begin at the last post-water touch, so the two are matched in what precedes them (consumption) and diverge only at the end (**Extended Data Fig. 12b**). Across 17 sessions, 1,194 trials had a post-water touch; of these 139 ended in an anticipatory hold and 585 in a terminal replace completed within 0.5 s (19.2% and 80.8% of the 724 analyzed; **Methods**). The hold is an anticipatory movement, and a postponement of the terminal replace. When timed from the last touch (consumption phase), a terminal replace follows a hold significantly (median 1 s) later than on replace trials, where it follows consumption immediately (**Extended Data Fig. 12c**). Each ending occurs on its own trials, which lets us ask whether ET^d^ and ET^i^ track the proactive and reactive endings in the same cells or in different ones.

For each population we took the 0.5 s following each ending’s own onset and asked whether the activity pattern there could be distinguished from the 0.5 s immediately preceding it, against a shuffle null; the two windows are identical in width and placement for the two endings and only the anchor differs (**Extended Data Fig. 12b**). Because both endings begin as consumption ends, the preceding window carries the same behavioral content for each, so the two are measured against the same reference, and each ending is tested only on its own trials against its own null, making the two measurements independent. As whole populations, both ET^d^ and ET^i^ decode both endings. ET^d^ reached 0.638 for the hold and 0.677 for the replace, and ET^i^ 0.817 and 0.821, against nulls at 0.50 throughout (**Extended Data Fig. 12d**). Cell by cell, among cells with at least five trials of each ending (1,193 ET^i^ and 110 ET^d^), 126 ET^i^ cells (10.6%) decoded the hold alone against 2.4% expected by chance (binomial p < 10^−40^), 41 (3.4%) decoded the replace alone (chance 2.1%; p = 0.002) and 18 (1.5%) decoded both (chance 0.1%; **Extended Data Fig. 12e,f**). Most hold-only ET^i^ cells decreased their activity at hold onset (79 of 126; **Extended Data Fig. 12g**). In ET^d^, 6 cells (5.5%; chance 2.5%; p = 0.06) decoded the hold alone, 5 (4.5%; chance 1.3%; p = 0.015) the replace alone and 1 both; with only 12 such ET^d^ cells, the mix of classes did not differ between ET^d^ and ET^i^ (χ^2^ p = 0.3). ET^i^, which as a whole decodes both endings, therefore contains a dedicated proactive-hold channel, a smaller dedicated reactive-replace channel and cells that decode both, reflecting another element of functional diversification.

## Discussion

Our results establish neurogenic origin as a developmental axis that configures the functional organization of cortical output. Rather than partitioning L5 ET neurons into separate transcriptomic classes, dNG and iNG generate two populations that span across transcriptomic ET types, yet diverge significantly through graded, coordinated shifts in effector-gene families, connectivity molecules, ion channels, receptors, synaptic organizers, axon-guidance proteins and transcription factors. This signature is not restricted to L5 ET, with L6 CT and L5 NP also showing the same iNG-biased effector gene shifts, and every DEG shared between subclasses changing in the same direction. The ET^d^ behave as a compact molecular and functional subpopulation with few distinguishing effectors and form a small, high-dimensional pool of heterogeneous response types that is generic to the behavioral task. The ET^i^ diversify these and elaborate it into a larger, specialized repertoire, at the molecular level through expanded effector-gene programs and, functionally, into low-dimensional, spatially clustered channels that track successive phases of the RGD behavioral sequence, extend coding to spontaneous movement, and separate proactive from reactive RGD terminal actions.

A major implication of our data is that neurogenic origin is a plausible developmental axis that generates cortical neuron functional heterogeneity. Indirect neurogenesis is the developmental route that generates within-subclass effector-gene variability in the adult cortex. ET^d^ and ET^i^ jointly populate every deep-layer ET transcriptomic type and share the core L5 ET corticofugal identity program^8,9,21^. Thus, every classic transcriptomic type is generated by both neurogenic mechanisms. They are distinguished by the graded, coordinated tuning of effector-gene families, including connectivity, channel, receptor and synaptic genes and this tuning is systematically patterned by neurogenic origin. Neurogenic origin thus defines a new layer of transcriptomic heterogeneity orthogonal to the classical transcriptomic types, organizing graded effector-gene variation within them. Because L6 CT and L5 NP neurons, which project to different targets, carry the same iNG-biased effector programs, this is likely a general feature of deep-layer PN^i^ subclasses rather than of ET^i^ alone. Indeed, recent work has suggested that within-subclass, graded transcriptomic variation, on top of defined discrete transcriptomic subclasses, is the hallmark of cortical and hippocampal neuronal heterogeneity, with continuous gene-expression gradients shaping much of the functional diversity within each type^74,75^. NDD-risk genes are likewise biased toward the iNG-derived population in all three PN subclasses, suggesting that indirect neurogenesis may be a shared point of vulnerability of deep-layer output neurons in neurodevelopmental disease. This framework also carries an evolutionary implication. The massive expansion of iNG in mammalian cortical evolution^18,76^ offers a powerful means to generate the vast number of functionally distinct output channels that the cortex requires, and our data suggest that graded effector-gene variability is a plausible mechanism through which this diversity may be achieved. By tuning a conserved corticofugal program rather than inventing new cell classes, iNG can efficiently scale functional diversity.

Whereas the transcriptomic diversification we describe is a general property of ET neurons across the isocortex and extends to L6 CT and L5 NP neurons, its functional consequences are most directly read out in the motor areas, where ET output drives movement^1,3,5,77^. Focusing on MOs, we find a functional counterpart to the molecular diversification which shapes how iNG contributes to cortical motor control. The cortex orchestrates skilled forelimb behavior by distributing control of successive movement phases across its ET neurons^77–79^. We find that both ET^d^ and ET^i^ encode movement kinematics, the common ’currency’ of motor-cortical activity^67,71,72^, providing a shared functional platform. On top of this, the ET^i^ resolve into behavior-epoch-specific channels that track distinct phases of the RGD sequence, and our action-selection model^69,70,80^ shows that this organization is sufficient to ’orchestrate’ it. Importantly, this functional diversification is itself graded and spatially structured. ET^i^ cells sharing an epoch preference lie near one another. Just as the generic effector-gene variation across cortical ET neurons is continuous^74,75^, the functional channels it specifies are graded, carried by single cells within a continuum rather than by discrete types. Thus, the graded effector-gene variability that defines a new transcriptomic layer, similarly manifests functionally, as a new layer of sequenced motor control. Further, ET^i^ diversification extends beyond movement kinematics to a higher level of control, generating dedicated channels that distinguish proactive anticipatory from reactive terminal actions^81,82^. Critically, the proactive and reactive endings are not merely different movements, but reflect distinct internal states, with a reactive ending executing the stereotyped close of the sequence, while a proactive ending expresses an anticipatory, goal-directed expectation of continued reward^81–85^. In keeping with the established role of MOs in representing such internal and anticipatory variables, from preparatory activity that predicts an upcoming action, to the decision of whether and when to act^73,81–83^, the dedicated ET^i^ channels for the two endings encode a cognitive variable, the animal’s predictive state, along with motor output^84–86^. Additionally, it is pertinent to note that these functional distinctions map onto anatomical differences. The same iNG program massively expands and dominates ET^i^ projections^15^ to brainstem action diversification^87,88^ and spinal action execution^3^ centers, so that the fine-grained, phase-specific functional channels built by ET^i^ are the activity counterpart of its elaborated descending connectivity. Together, these results position ET^i^ as ’*diversifiers*’ that refine a generic ET^d^ action representation, into specific, sequenced control of skilled behavior, with iNG supplying both the molecular and the functional substrate for this expansion. Since L6 CT and L5 NP neurons carry the same iNG-biased effector programs, it will be important to test whether indirect neurogenesis similarly diversifies their output, for example corticothalamic feedback from L6 CT, into specialized channels.

Another consideration concerns how this diversification scales with behavioral dexterity over evolution^5,89^. In the primate, skilled hand control emerges from the cooperative division of labor between two cortical descending systems, the corticospinal pathway providing fine, individuated muscle control and the corticofugal pathway providing broad, synergistic control^90,91^, a specialization that accompanies the expanded behavioral repertoire of the primate hand^1,2,92^. Indirect neurogenesis in the primate, may be scaled to build the enlarged and functionally specialized descending machinery that underlies fine digit control^2,93–95^. Thus, the graded effector-gene diversification by indirect neurogenesis is plausibly a developmental substrate that evolution can expand, through the output pathways, to meet escalating demands for behavioral precision^2,5,95,96^.

A question that follows naturally is, what does direct neurogenesis contribute? Our data motivate a unifying framework in which ET^d^ and ET^i^ embody two complementary developmental strategies. Transcriptomically, this small ET^d^ pool carries few physiological and connectomic distinguishing effectors while still contributing to all transcriptomic cell types. Functionally in MOs, a small, high-dimensional ET^d^ pool spans a broad, heterogeneous range of response types that is generic to the RGD task, with higher spontaneous activity and a more dispersed response-profile space than ET^i^. Indirect neurogenesis acts on this substrate, reproducing and amplifying the templates and, through graded effector-gene tuning, sharpening its broad, spontaneously active repertoire into the specific, epoch-tuned channels that structure sequential behavior. Our earlier optogenetic activation of ET^d^ and ET^i^ in the motor cortex (rostral forelimb orofacial area, RFO^97^) showed that ET^d^ generate limited and small head/orienting movements whereas ET^i^ not only amplify the head/orienting movements but additionally generate forelimb and orofacial movements, essentially the full repertoire of behavior as elicited by activation of the pan-FezF2 population^15^. This ET^d^ role recalls a conceptual parallel in cortical development, in which an early, ancestral population lays down a template that later, expanded populations build upon. Pioneer neurons, the earliest-born subplate cells, establish the first axonal scaffolds and transient circuits that guide the later-arriving cohorts which come to dominate the mature cortex^98,99^. It is also important to note here, that we have demonstrated that axonal refinement via pruning for target specificity during postnatal development occurs exclusively in ET^i^, following an exuberant axonal overgrowth phase shared with the template ET^d^^15^. ET^d^ thus occupy an analogous position along the neurogenic axis, as the population that provides the generic corticofugal template, which we deem as ’*archetypes*’, which indirect neurogenesis subsequently expands and diversifies. In the context of evolution, because dNG is the ancestral and more conserved route, the ET^d^ archetype may represent a transcriptomically and functionally conserved core, shared broadly across species, whereas the iNG diversifiers, may carry the species-specific functional classes that distinguish them. The transcriptomic data suggest that this division of labor is shared also by the other deep-layer PN subclasses. By defining these complementary contributions, our work assigns direct and indirect neurogenesis distinct and definable roles in constructing the functional mosaic of cortical cell types. Direct neurogenesis builds generic archetypes, and indirect neurogenesis elaborates it into the specific, expandable functional classes through which the cortex may scale cell types mediating the diversity and precision of behavior.

## Methods Mice

All animal procedures conformed to National Institutes of Health guidelines and were approved by the Institutional Animal Care and Use Committee of Duke University. Adult male and female mice (8 -16 weeks at the start of imaging) were used and maintained on a 12-h light/dark cycle with food and water available *ad libitum* except during the water-restriction period required for the behavioral task.

Deep layer *Fezf2* expressing projection neurons (PN) across the isocortex generated through Direct-(PN^d^) and indirect-(PN^i^) neurogenesis were distinguished using the *Fezf2TC;Tbr2-Flp* double-knock-in mice^15^. The deep-layer PN consisted of the following subclasses^7^: L5 ET (extratelencephalic neurons),L5 NP (near-projecting neurons) L6 CT (cortico-thalamic neurons) and L6b neurons. For 2P-imaging, a tTA-dependent (TRE) AAV delivered the red indicator jRGECO1a to ET^d^ neurons, and a Cre-dependent (FLEX/DIO) AAV delivered the green indicator GCaMP8f to ET^i^ neurons, co-injected into secondary motor cortex (MOs) so that both subtypes were labelled and imaged simultaneously within a single field of view in Layer 5b. For the subtype-blind reference, the pan-ET population was labelled in a separate cohort by crossing Fezf2-CreER mice to a jGCaMP8m reporter line (*TIGRE2-jGCaMP8m-IRES-tTA2-WPRE*), so that CreER-mediated recombination drove GCaMP8m across the entire ET class without distinguishing neurogenic origin. For single-nucleus RNA sequencing, PN^d^ and PN^i^ nuclei were labelled in *Fezf2-CreER;Tbr2-Flp;IS-INTACT*^13,15,100^ mice where PN^d^ nuclei are RFP^+^ and PN^i^ nuclei GFP^+^ (**Fig. 1a,b**). For the *IS-INTACT* mice, briefly, a STOP cassette flanked with *loxP* sites and dTomato (dimer) sequence flanked with *FRT* sites was targeted at the *Gt(ROSA)26Sor* locus, preventing the transcription of enhanced green fluorescent protein (EGFP). dTomato is expressed in the nuclei following *cre*-mediated recombination, while EGFP is expressed in the nuclei following both *cre*-and *flp*-mediated recombination.

## Tamoxifen induction

Tamoxifen (T5648, Sigma) was dissolved in corn oil (20 mg/ml) with constant magnetic stirring overnight at 37 °C. For *Fezf2TC;Tbr2-Flp* mice carrying viral infections, tamoxifen was delivered by intraperitoneal injection at 100 mg/kg, with the first induction one day after viral infection and subsequently every 2–3 days for 2–3 doses, to drive CreER-dependent recombination and thereby restrict the Cre-dependent indicator to the ET^i^ subtype. A similar dosage was administered to *Fezf2-CreER;Tbr2-Flp;IS-INTACT* mice at P21 and P28. The subtype-blind pan-ET cohort (*Fezf2-CreER;jGCaMP8m* reporter mice) received tamoxifen at the same concentration and dose to induce CreER-dependent expression of GCaMP8m across the ET class prior to imaging.

### Single-nucleus RNA sequencing Sample acquisition

*Fezf2-CreER;Tbr2-Flp;IS-INTACT* triple transgenic animals were generated by crossing *Fezf2-CreER*^13,20^ mice (Jackson Laboratory stock no. 036296) with *Tbr2-Flp*^13^ and *IS-INTACT* mice and maintained on a C57BL/6 background. At P30-P35, mice were anesthetized with isoflurane and decapitated. Brains were rapidly removed and put into ice cold Hibernate A supplemented with B27 and Glutamax for microdissection for the whole neocortex. Dissected tissue was flash frozen on dry ice with isopentane within 15 min of decapitation and stored at -80 °C until use.

### Nuclei sorting

Nuclei were isolated using the Nuclei EZ Prep kit (Sigma Aldrich, NUC101). Frozen tissue was homogenized in ice-cold lysis buffer, and nuclei were recovered by centrifugation at 500 × g at 4 °C. Isolated nuclei were resuspended in nuclei suspension buffer (1x PBS, RNAse inhibitor, BSA), filtered through a 40-µm strainer, and stained with Vybrant DyeCycle prior to FACS. GFP-positive and RFP-positive nuclei were isolated by fluorescence-activated sorting using a Beckman Coulter sorter with a 70-µm nozzle. Nuclei were gated by size and Vybrant dye signal before sorting the GFP-positive and RFP-positive populations into separate collection tubes.

### Library construction and sequencing

Sorted nuclei were processed using the Chromium Next GEM Single Cell 3′ Reagent Kit v3.1 (10x Genomics) according to the manufacturer’s protocol. Libraries were prepared for single-nucleus RNA sequencing and sequenced by the DNBSEQ-G400/T7 platform.

### Sequencing data processing and quality control

Sequencing reads were aligned with Cell Ranger 7.0.0 (cellranger count, --include-introns=true) to the mouse reference transcriptome (pre-built Cell Ranger reference package mm10-2020-A) extended with the reporter sequence (EGFP for ET^i^, mRuby for ET^d^ libraries). Reporter features were removed from the count matrices before analysis. A QC on the twelve libraries was performed. First, ambient RNA was estimated per library with SoupX^101^. Then, libraries were excluded if they were flagged on two or more of four measures: SoupX contamination, fraction of reads in cells, median genes per nucleus and median UMIs per nucleus. A library was flagged on a measure if its value lay more than 1.5 × the interquartile range beyond the quartile in the unfavorable direction, across all 12 libraries. Based on this rule, two batch-2 libraries were excluded (one PN^i^, one PN^d^), leaving six PN^i^ and four PN^d^ libraries. Within each library, nuclei with 2,000–70,000 UMIs, more than 1,000 detected genes and less than 5% mitochondrial UMIs were retained, and doublets were removed with scDblFinder^102^ (default settings). This left 92,830 nuclei. Count matrices were merged and analyzed in R (v4.6.1) with Seurat (v5.5.1)^103^. Counts were log-normalized jointly across all nuclei (scale factor 10,000), and the 2,000 most variable genes were scaled across all nuclei together. Principal components were integrated with Harmony^104^ on sequencing batch, each batch containing both PN^d^ and PN^i^. Nuclei were annotated with the MapMyCells^105^ web portal (RRID:SCR_024672) from the Allen Brain Institute against the 10x Whole Mouse Brain taxonomy (CCN20230722), by hierarchical mapping on raw counts. Analyses used glutamatergic nuclei of the four deep-layer PN subclasses targeted by *Fezf2* (L5 ET, L6 CT, L5 NP and L6b) whose class and subclass bootstrapping probabilities both exceeded 0.9. Analyses at cluster level also required supertype and cluster probabilities above 0.7. Ninety genes were removed from all analyses and enrichment backgrounds: 83 markers of non-neuronal cells (vascular, glial, immune, meningeal and blood; **Supplementary Table 3**), and 7 genes (Klf2, Ramp2, Ctla2a, Adgrl4, Aldoc, Rplp1, Tmsb4x) whose dNG/iNG difference in a preliminary analysis was explained by the SoupX ambient profiles.

### Comparison of cell-type landscape between dNG and iNG

Cell-type landscape was compared between dNG and iNG in nuclei with class and subclass bootstrapping probabilities above 0.9 and supertype and cluster probabilities above 0.7, using MapMyCells cluster labels as cell types. The UMAP was recomputed for this population from the first 30 Harmony components.

#### Depth matching

Where PN^d^ and PN^i^ nuclei were compared directly, they were matched within each cluster: nuclei were binned by number of detected genes (deciles), and equal numbers of PN^d^ and PN^i^ nuclei were drawn from each bin. Clusters with fewer than 25 nuclei of either population were dropped, leaving 29,138 nuclei in 57 clusters.

#### Pairwise differences between cell types

The number of differentially expressed genes was counted between every pair of clusters, separately within PN^d^ and within PN^i^ nuclei of the depth-matched set^106^. Each pair was subsampled to equal size and tested with the Wilcoxon rank-sum test; a gene was counted if adjusted p < 0.05, the difference in mean log-normalized expression exceeded 1, and it was detected in at least 10% of nuclei of one cluster. For each cluster, the 0.1–0.9 quantiles of its counts against all other clusters were taken, and the distributions of these quantiles were compared between PN^d^ and PN^i^.

#### De novo clustering

To ask whether either PN^d^ and PN^i^ resolves into more transcriptomic clusters without using the MapMyCells reference cell type labels, an PN^d^ set (all four PN^d^ libraries) and an PN^i^ set (all six PN^i^ libraries) were drawn from deep-layer PN nuclei (L5 ET, L6 CT, L6b and L5 NP together) with equal numbers of nuclei, matched in cluster composition and in detected genes. Clusters with fewer than 5 nuclei in either PN^d^ and PN^i^ (12 of 81 clusters, 0.4% of nuclei; none of them was present in only one neurogenic population) were excluded before the draw. Each set was clustered on its own on the first 30 Harmony components (shared nearest-neighbor graph, Louvain) at resolutions 0.2–3.0 in steps of 0.2. A cluster was counted as stable if its mean best-match Jaccard index across five reclusterings of random 80% subsamples was at least 0.6. The draw was repeated five times. As a null, each draw’s nuclei were pooled and split into two mixed sets with the same matching, ignoring their neurogenic origin (15 splits).

#### Cluster correspondence

Stable clusters with at least 30 nuclei from the first draw were matched between PN^d^ and PN^i^ at each resolution by three measures of similarity. (1) *Expression*: each cluster was summarized by its mean log-normalized expression of the variable genes, centered across all clusters of both ET^d^ and ET^i^, and the similarity of an PN^d^ and an PN^i^ cluster was the Pearson correlation of these profiles. (2) *Cell-type composition*: each cluster was described by the fraction of its nuclei in each MapMyCells reference cluster, and similarity was the cosine similarity of these fractions. (3) *Marker overlap*: marker genes of each cluster were identified within PN^d^ and PN^i^ by a Wilcoxon test against the other clusters (adjusted p < 0.05, log fold change > 0.25), the top 100 by AUC were kept, and similarity was the Jaccard index of two clusters’ marker lists. A match required a similarity above the 95th percentile of a null distribution. For expression, the null was the best-match similarities after randomly relabeling PN^i^ nuclei (50 permutations), and the same threshold was used for composition. For marker overlap, the null was the best-match Jaccard indices between random gene lists of the same sizes drawn from all markers found (50 permutations). Each PN^i^ cluster was linked to its most similar PN^d^ cluster, using each similarity measure separately.

### Differential expression and gene-family enrichment

We performed differential expression analysis between PN^i^ and PN^d^ within each subclass on pseudobulk profiles, with libraries as replicates. Mitochondrial genes and predicted genes were removed, as were the sex-marker genes Xist, Tsix, Ddx3y, Eif2s3y, Kdm5d and Uty. Models were fitted with DESeq2^107^ using the design ∼batch + neurogenic origin. Sex was not included as a covariate . Its effect was tested directly by sex balancing (below). A gene was called significantly differentially expressed (DEG) at adjusted p < 0.05, |log2 fold change| ≥ 0.25 and detection in at least 10% of the subclass’s nuclei in either neurogenic origin.

Gene families were tested by gene set enrichment analysis (GSEA^108^). Within each subclass, all tested genes were ranked by the DESeq2 Wald statistic for the dNG/iNG contrast, and enrichment was computed with clusterProfiler^109^ for ∼2,000 HGNC (HUGO Gene Nomenclature Committee) gene families together with 38 gene families (see^110^) with Benjamini–Hochberg adjustment. A positive normalized enrichment score (NES) indicates a family shifted toward higher expression in PN^d^. The analysis was repeated within batch 1 alone and within batch 2 alone. Gene families with adjusted p < 0.05 and DEGs in the analysis of both batches were then subjected to two checks, applied in the same way to both.

#### Batch concordance

A family or DEG was retained if its NES or log2 fold change had the same sign in each batch analyzed alone.

#### Sex balancing

Nuclei were called female (at least one Xist UMI and no UMI from the Y-linked genes Ddx3y, Eif2s3y, Kdm5d and Uty) or male (at least one Y-linked UMI and no Xist UMI); others were left unassigned. Within each batch, female nuclei were removed at random from each ET^d^ library whose female fraction exceeded that of the pooled ET^i^ libraries, until the two matched. The DESeq2 model and GSEA were rerun on the balanced data for five independent random draws, and a result passed if it remained at adjusted p < 0.05 with the same sign in every draw.

#### Member genes

For each reported family, each member gene was described on log2(DESeq2-normalized pseudobulk + 1). A member was called consistently shifted if its PN^d^-PN^i^ difference had the same sign in batch 1 and batch 2, both before and after removing the batch effect (limma removeBatchEffect^111^); at least 75% of PN^d^–PN^i^ library pairs differed in the same direction as the model fold change after batch removal; |log2 fold change| was at least 0.25; and it was detected in at least 10% of the subclass’s nuclei in either PN^d^ and PN^i^. The same criteria were required to hold in every sex-balanced draw for a gene to be called consistently shifted in all analyses.

### PN^d^ and PN^i^ classification by gene-family expression

Whether the gene families identified by GSEA could also distinguish individual PN^d^ and PN^i^ nuclei was tested with MetaNeighbor^112^. Within each subclass, PN^d^ and PN^i^ was set as the label to be discriminated and sequencing batch as the study label. Each AUROC measures how well a family’s expression, learned from the nuclei of one batch, separates PN^d^ and PN^i^ in the other; 0.5 indicates no separation. MetaNeighbor was run on raw counts of the analysis population for all gene families. AUROCs were converted to p values with the normal approximation to the Mann–Whitney U statistic and adjusted by Benjamini–Hochberg within each subclass. A family was considered to classify PN^d^ and PN^i^ above chance if it had an FDR < 0.05, an AUROC of at least 0.6, and at least five member genes detected in at least 10% of the subclass’s nuclei.

### Single-gene differential expression within gene families

Gene-family panels (**Fig. 2c-h; Extended Data Figs. 3, 4**) show only DEGs. In L5 ET, the genes shown are the consistently shifted members of the enriched families (see Member genes) that are also DEGs, grouped into six functional categories: ion channels and Ca^2+^ transporters; transcription factors; synaptic organizers, scaffolds and Ca^2+^/CaM signalling; Ig/I-set cell-adhesion molecules and cadherins; axon guidance, adhesion and cytoskeleton; and modulatory and peptide receptors. For L6 CT and L5 NP, the DEGs of each class were assigned to the same six categories by manual curation of gene function and HGNC family membership.

### Cross-subclass concordance of DEGs

For each pair of subclasses (L5 ET and L6 CT, L5 ET and L5 NP, L6 CT and L5 NP), the genes that were DEGs in both subclasses were compared by the sign of their log_2_FC (**Fig. 2b**). Agreement was tested with a two-sided exact binomial test against 0.5.

### Neurodevelopmental-disorder risk genes

The DEGs of L5 ET, L6 CT and L5 NP were matched to human neurodevelopmental-disorder (NDD) gene resources: SFARI Gene^57^ (release of 28 March 2024), Genes4Epilepsy^113^ (v2026-09), the exome-wide significant genes^114^, GenCC gene–disease validity submissions (snapshot of 24 December 2021), the SCHEMA exome-wide significant schizophrenia genes^115^ and PsychENCODE^116^ high-confidence schizophrenia genes; DEGs not covered by these resources were checked in OMIM^117^ and the primary literature. Each gene was assigned to one of three tiers of human genetic evidence: strong (SFARI score 1 or syndromic; Genes4Epilepsy; DDD; GenCC definitive or strong; published Mendelian NDD), limited (SFARI score 2; GenCC moderate; a single family or few cases; replicated or meta-analysed association) or weak (SFARI score 3; GWAS-mapped only; single unreplicated association; no human evidence). Genes with strong or limited evidence and a neuronal function (synaptic, channel, receptor, adhesion, guidance or neuronal signaling, as opposed to housekeeping, metabolic or organelle genes; assigned by curation) were retained as NDD-risk genes and assigned to epilepsy, autism, intellectual disability/developmental delay or schizophrenia/psychiatric categories by their principal association.

### Two-photon calcium imaging during skilled reaching Stereotaxic surgery

Mice were anaesthetized with isoflurane (2-5% induction, 0.8-1.2% maintenance) and positioned in a stereotaxic frame, with body temperature maintained at 34–37 °C. Lidocaine (2%) was applied subcutaneously to the scalp and ketoprofen (5 mg/kg) administered intraperitoneally before and after surgery. A midline incision exposed the skull, a burr hole was drilled over secondary motor cortex, and a pulled glass pipette (tip 20–30 µm) loaded with viral suspension was lowered to the target. MOs was targeted at +1.6 mm anteroposterior, 1.37 mm mediolateral and −0.7 mm dorsoventral relative to bregma. A volume of 400–700 nl was infused at 10–30 nl/min with a Picospritzer (General Valve), and the pipette held for 5 min before slow retraction to prevent backflow; the incision was closed with tissue adhesive (3M Vetbond) and nylon suture and mice recovered on a heating pad.

### Head-fixed reach-grasp-drink behavior

The behavioral paradigm was a head-fixed reach-grasp-drink (RGD) task. Water-restricted mice were head-fixed and trained to reach with the left forelimb for a water droplet presented at a spout, withdraw the hand to the mouth, and drink by licking water from the hand. Mice were trained until performance stabilized (>80% completed trials) before imaging. The spout was presented at one of three positions (P1-P3), randomized across trials. The dual-colour dataset comprised 17 sessions from 4 animals and 1,684 trials in total (≈100 per session); the pan-ET cohort comprised 10 sessions from 3 animals and 900 trials. Trials were 475 imaging frames long (≈15.9 s at the acquisition frame rate). Starting from the last post-water spout touch, Replace onset was the first frame at which the hand’s distance from the spout exceeded 50% of that trial’s reach extent (the pre-lift-to-spout distance), a relative criterion that scales to each trial’s geometry. Every post-water-touch trial was classified either as return (hand retreated past the threshold, a completed reach, drink and withdraw cycle; 972 trials) or hold (hand remained near the spout, staying beyond the threshold for ≥85% of a ≥1 s post-touch window, 139 trials). A further 490 trials had no post-water touch anchor and thus excluded from Replace analyses.

### Two-photon calcium imaging

Two-photon imaging of MOs layer-5 ET was performed in head-fixed mice during behavior. In dual-colour sessions, ET^d^ (jRGECO1a^65^, red) and ET^i^ (GCaMP8f^66^, green) neurons were imaged simultaneously within a single field of view, with the two indicators separated into red and green detection channels, at a frame rate of 30 Hz (475 frames per trial). In separate pan-ET sessions the entire ET population was imaged in a single indicator (GCaMP8m^66^; ET^all^).

### Behavioral tracking and kinematics

Forelimb and orofacial movements were captured with two synchronized high-speed cameras (Teledyne FLIR) fitted with varifocal lenses, recording the mouse from the front and the side under infrared illumination at 150 frames per second and providing calibrated views for three-dimensional reconstruction; the geometric relationship between cameras was obtained with the MATLAB Camera Calibrator. Two deep neural networks, one per view, were trained in DeepLabCut 3.0.0^118^ (PyTorch engine, ResNet-50 backbone) to track keypoints on the digits, hand, nose, mouth, tongue, spout and water drop (18 keypoints in the front view and 22 in the side view), using manually labelled frames sampled across animals, body sizes and illumination conditions. Predicted points below a confidence of 0.6 were treated as missing. Three-dimensional hand and spout positions were reconstructed by stereo triangulation. Forelimb and orofacial kinematics were extracted from high-speed video by marker pose estimation and converted to per-frame series aligned to the imaging clock. The continuous series entered into the encoding model were: three-dimensional hand position (x, y, z), three-dimensional hand speed and cumulative path length, vertical speed, aiming score, digit aperture, hand-to-spout and hand-to-mouth distance, nose displacement, mouth aperture, right-hand speed, water area, spout position (x, y), tongue length and jaw displacement.

### Image processing and cell selection

Data were acquired on a Bruker Prairie two-photon microscope. Motion correction, ROI detection and fluorescence extraction were performed in Suite2p^119^. Indicator-appropriate decay constants were used (jRGECO1a τ ≈ 1.25 s; GCaMP8f τ ≈ 0.4 s; and, for the separately recorded pan-ET, GCaMP8m at τ ≈ 0.4 s). For every dual-colour session we computed the pixel-wise intersection-over-union (IoU) between each accepted ROI in one channel and each accepted ROI in the other, and flagged pairs with IoU ≥ 0.3 as the same physical cell. Such ROIs were removed from *both* channels. Sessions lacking a valid strict mask were rejected rather than falling back to raw Suite2p labels. This correction reduced the accepted populations, from 684 to 197 ET^d^ and from 2,282 to 1,816 ET^i^ neurons across 17 sessions. dF/F_0_ was computed against a static per-cell baseline defined as the whole-recording mean of the neuropil-corrected trace, without detrending, and all analyses used dF/F_0_ unless stated otherwise. Deconvolved activity estimates from Suite2p (OASIS) were used as a control for indicator kinetics in the single-cell peri-epoch decoding (**Extended Data Fig. 8b**) and as the input to the basal-ganglia model. Spike inference for the single-indicator ET^all^ used CASCADE (see Indicator controls).

### Definition of action phases and peri-event epochs

Action landmarks (water delivery, lift onset, aim completion, advance endpoint, grasp completion, touch, supination, first lick, replace onset) were taken from the behavioral annotation. Four peri-event epochs were defined as windows from 0.5 s before to 1.0 s after an anchoring event: peri-spout (spout arrival), peri-water (the first reach after water delivery, detected as lift onset), peri-consume (the first spout touch after water delivery) and peri-replace (replace onset, on returned trials). Their boundaries were set by the measured spacing of the landmark events against the decay kinetics of the red indicator since landmarks separated by less than the decay floor of ≈0.5 s cannot be resolved as independent calcium transients. An epoch contributed on a given trial only if its anchor lay at least 1.5 s (the window width) from every other anchor on that trial, so that windows never overlapped, and peri-replace additionally required replace onset to follow the first lick by at least 0.5 s, to avoid contamination from the preceding consummatory transient. Of 1,684 trials, peri-spout was therefore usable on all, peri-water on 645, peri-replace on 463 and peri-consume on 173 (**Extended Data Fig. 8a**). Single-cell decoding used all four epochs; the single-cell encoding model used the three epochs up to consumption (peri-spout, peri-water and peri-consume; Fig. 4c), and the basal-ganglia model the three behavioral epochs (peri-water, peri-consume and peri-replace).

### Spontaneous transients in task-free windows

Task-free movement (TFM) windows were defined per trial as the pre-spout and post-replace periods, which contain movement but no task demand, with a minimum duration of 1 s. Spontaneous transients were detected on the corrected dF/F_0_ traces as excursions past a fixed threshold of ±0.15, of either sign, and peak-aligned raw waveforms were clustered into eight transient families. Event rate was computed per cell as events per second within the window type.

Because the populations differ greatly in cell number, across-population comparisons subsampled each population to the ET^d^ cell number (n = 197) over 50 draws and were assessed by a label-permutation test passed through the identical matched-subsampling estimator, Bonferroni-corrected for the three pairwise comparisons. The same comparison was made in the RGD window (**Extended Data Fig. 6g,h**). Within-population comparisons of task-free versus task windows were paired across cells (Wilcoxon signed-rank).

### Whole-trial response profiles

For each cell a whole-trial response profile was formed as the trial-averaged dF/F_0_ across the 475-frame trial, baseline-subtracted using the first 50 frames, lightly smoothed (Gaussian, σ = 1 frame) and z-scored.

These profiles are the feature space for the population trajectories, effective dimensionality and dispersion analyses below. Ordered cell–cell correlation matrices (**Fig. 4b**) show the Pearson correlations between these profiles, with cells ordered along the dominant response axis.

### Population trajectories and divergence

Population trajectories were constructed by binning each cell’s trial-averaged response into 20 equal time bins across the trial and reducing the resulting bins × cells matrix to three principal components, giving a 20-point path through state space per population. Trajectories were compared after Procrustes alignment and divergence is reported as the Procrustes disparity, and as the per-bin Euclidean distance along the trial. Comparisons between populations were made at matched cell count (n = 197, the ET^d^ number; 200 draws), and significance was read from the label-permutation null (200 permutations; P = 0.005 is the smallest attainable value; **Extended Data Fig. 6e**).

### Effective dimensionality

The effective dimensionality of a population’s response manifold was quantified as the participation ratio, PR = (Σ_i_λ_i_)^2^/Σ λ ^2^, where λ are the eigenvalues of the covariance matrix of the cells’ whole-trial response profiles. ET^d^ and ET^i^ were compared at matched cell count (n = 197) over bootstrap draws, reported with 95% intervals and as the fraction of draws in which ET^d^ exceeded ET^i^ (**Fig. 3h**). Within ET^all^, the same measure was computed for cells matched cell by cell to ET^d^ or to ET^i^ at matched n = 40, with significance from the reference-shuffle permutation described there (**Extended Data Fig. 6i**).

### Cell-to-cell correspondence between ET^d^ and ET^i^

To relate the response repertoires of the two populations without clustering, each ET^i^ cell was compared with every ET^d^ cell by the Pearson correlation of their trial-averaged dF/F_0_ profiles over the RGD window (−2 to +10 s from spout onset) and assigned to the ET^d^ cell with the highest correlation. A match was counted when this correlation exceeded r = 0.79, the 95th percentile of a null of best-match correlations after circular time-shifts of the profiles. For each ET^d^ cell, the number of matched ET^i^ cells (fan-in) was counted, and ET^d^ cells were classed as receiving one ET^i^ cell (one to one), several (one to many) or none (**Extended Data Fig. 7a**).

### Dispersion in response-profile space

The dispersion of the population was the mean pairwise Euclidean distance between the z-scored whole-trial response profiles of its cells. ET^d^ and ET^i^ were compared under a matched-n bootstrap at the ET^d^ cell number (1,000 draws), reported as the fraction of draws in which one exceeded the other and as the 95% interval of the difference (**Extended Data Fig. 7b,c**). Its dependence on sample size was mapped by drawing 1,000 random subsets of each size from each population (**Extended Data Fig. 7f,g**).

### Indicator controls using the single-indicator pan-ET population

ET^d^ and ET^i^ are reported by different indicators while the pan-ET recordings are a separate cohort of animals imaged with a single indicator (GCaMP8m). The following control analyses were performed:

#### Composition

Each ET^all^ cell was assigned to whichever single ET^d^ or ET^i^ cell its whole-trial profile correlated with most strongly. Cells were thereby assigned on response profile alone and it was compared with the 9.8% ET^d^ measured in the dual-color imaging by a binomial test (**Extended Data Fig. 7d**).

#### Dispersion and dimensionality

Dispersion and participation ratio of the ET^d^-matched and ET^i^-matched ET^all^ cells were compared at a common size of 40 cells per draw, strictly below the size of the smaller group so that each draw has genuine sampling variability. The null shuffles which reference cells count as ET^d^ and which as ET^i^, holding their numbers fixed, and repeats the whole matching, thereby asking whether the difference depends on the reference cells being the actual populations (**Extended Data Figs. 6i, 7e**).

#### Spike inference

To test whether the peri-epoch structure is a property of calcium dynamics rather than of spiking, spike rates were inferred from the pan-ET recordings with CASCADE^68^ model using the GCaMP8m-specific model at 30 Hz with 25 ms ground-truth smoothing, matching the acquisition rate of 30 Hz. Traces were converted from the pipeline’s mean-referenced dF/F_0_ to the percentile-referenced, percent-scaled form the models are trained on; inferred rates averaged 5.5 Hz, consistent with layer-5 ET neurons during task engagement. The single-cell peri-epoch decoding (below) was then run unchanged on the inferred rates (**Extended Data Fig. 8b**).

### Single-cell peri-epoch decoding

For each cell and trial, dF/F_0_ was z-scored within the trial and averaged over each usable epoch window, giving one value per cell, epoch and trial. Epoch identity was decoded from this single value by leave-one-trial-out cross-validation, with the epoch means learned from all other trials, and each held-out observation assigned to the epoch whose mean was nearest (one-dimensional nearest centroid), all observations of a trial being held out together. Accuracy was compared with a null of 1,000 surrogates in which the epoch labels were permuted within each trial, preserving each trial’s composition, and a cell was counted as decoding the epochs when its accuracy exceeded this null at P < 0.05; chance for this criterion is 5%. Proportions of decoding cells were compared between ET^d^ and ET^i^ by Fisher’s exact test. A cell’s preferred epoch was the epoch with the highest one-versus-rest area under the ROC curve computed from the held-out distances, and preferred-epoch distributions were compared between ET^d^ and ET^i^ by χ^2^ test (**Extended Data Fig. 8c**). The analysis was repeated unchanged on Suite2p-deconvolved activity for both ET^d^ and ET^i^ and, for ET^all^, on CASCADE-inferred spike rates (**Extended Data Fig. 8b**).

### Task engagement

To ask whether a cell was engaged by the task, its mean dF/F_0_ was computed in every RGD and every TFM window occurrence across trials, and the separability of the two sets was measured as the area under the ROC curve (AUROC). Significance was assessed by permuting the RGD and TFM labels within the cell (500 permutations), and a cell was counted as task-engaged when P < 0.05 and AUROC > 0.5, that is when it was more active during RGD than during TFM (**Fig. 4d,e**).

### Single-cell encoding models

Encoding was assessed per cell with a cross-validated Gaussian generalized linear model on dF/F_0_, using five-fold cross-validation across trials, mild L2 regularization (α = 1) to handle collinearity in the design, and a minimum of eight trials per cell. Model fit is reported as deviance explained (DE) on held-out data, both for the full model and, by leave-one-out refitting, as the change in deviance explained attributable to each regressor (ΔDE). The *kinematic* design comprised the 18 continuous behavioral series listed above, each entered at five lags spanning 0 to 0.5 s (sampled at approximately 0, 0.1, 0.2, 0.35 and 0.5 s) to keep the design matrix tractable. The *peri-epoch* design comprised one boxcar regressor for each of the three peri-event epochs up to consumption (peri-spout, peri-water and peri-consume), with the remainder of the trial as the reference level, and no lags; deviance explained therefore measures variance attributable to which epoch a frame belongs to, over a baseline-only null. The kinematic model was fitted over the whole trial in two divisions, the RGD task window and the task -free (TFM) window; the peri-epoch design defines its own frames and was fitted over the RGD task window only. Every cell was referenced to its own null distribution of 100 circular time-shift surrogates. A cell was counted as encoding only if its DE exceeded the 95th percentile of its own surrogate distribution and was itself positive. Under this criterion the expected rate by chance is at most 5%. Encoding rates are read against this 5% rate and checked for consistency across the 17 sessions and the four animals as independent units (**Extended Data Fig. 11g**); a binomial test against chance is used only where a single rate is tested directly, as for the task-free ET^d^ result. To separate task-driven from task-free movement encoding, the kinematic model was refitted on the same cells within the task-free windows, using the identical design, cross-validation scheme and surrogate null. Because the same cells are fitted in both windows, task and task-free encoding were compared within cell (Wilcoxon signed-rank on paired DE, and cross-tabulation of the binary encoding calls).

### Spatial organization of peri-epoch preference

Each imaging session is an independent field of view. Cells were positioned by their Suite2p ROI centroid in the imaging plane. All ET^i^ cells that decoded the four peri-epochs above their shuffle null on dF/F_0_ (721 of 1,816) contributed, each labelled by its preferred epoch. The statistic used was k-nearest-neighbor concordance: for each cell, the fraction of its k nearest other cells that prefer the same epoch, averaged over the cells of a field (k = 1, 5, 8 and 12; k = 12 for the main test). Each field was compared with a null of 2,000 permutations of the epoch labels among that field’s own cells. Per-field P values were combined across fields by Fisher’s method. To report the spatial scale, distances to the k-th neighbor were converted to micrometers. ET^d^ could not be tested with only 38 epoch-decoding ET^d^ cells across 17 fields.

### Basal-ganglia model of action selection

#### Model

Selection was simulated with the Gurney-Prescott-Redgrave (GPR) model of the basal ganglia in the form described previously^69^, used without modification. Each channel is represented by seven leaky-integrator units: striatal D1 and D2 populations, subthalamic nucleus, three globus-pallidus populations and the output nucleus (SNr/GPi), with piecewise-linear output functions and the published weights, thresholds, leak and timestep. The architecture implements a focused direct (D1) pathway that selectively inhibits the output nucleus and a diffuse subthalamic projection that excites it in proportion to the total salience across channels, so that a channel is released by disinhibition only when its own drive exceeds the summed drive of its competitors. Dopamine was fixed at 0.2 throughout. The full model, its parameters and its behavior across the salience plane are in^69^.

#### Circuit state

The circuit was run intact (diffuse = 1): the subthalamic projection to the output nucleus is the across-channel sum, giving the on-centre/off-surround competition.

#### Channels

The model was given three channels corresponding to the three behavioral peri-epochs, peri-water, peri-consume and peri-replace, presented in that order. Chance for correct selection is accordingly 1/3.

#### Salience

Channel salience was derived from deconvolved activity and epochs were event-anchored trial by trial using the windows, anchors and separability gate of the encoding analysis, so that trial-to-trial variability in event timing does not smear the response. For each ET^d^ and ET^i^, Pⱼ denotes the trial-averaged population response in window j and wₖ the mean of Pₖ over its own window, unit-normalised; the salience of channel k at time t during window j is the matched-filter output wₖ·Pⱼ(t). Diagonal terms therefore measure a channel’s response in its own epoch and off-diagonal terms its response during another epoch, so a channel that is active everywhere confers no advantage. The resulting drive was divided by a single global maximum to bring it into the 0-1 range the circuit’s thresholds assume.

#### Readout

A channel counts as selected when its SNr/GPi activity falls below half of its own tonic (pre-drive) level. Correct selection is the fraction of the driven timeline on which the most disinhibited channel is the one whose epoch is current; we also report the mean number of channels disinhibited per time bin, where one indicates clean single-channel selection.

### Proactive and reactive endings of the RGD action

Each trial with a post-water touch was classified from the kinematics as ending in a *hold*, the anticipatory proactive posture, or a *replace*, the reactive terminal movement. **Hold:** A trial counts as a hold when, after the last post-water touch, the hand remains near the mouth for at least 85% of a window of at least 1 s rather than returning to the perch. The hold is anchored on that last touch, which is the moment a return would otherwise have begun. **Replace:** A replace is anchored on the return itself, the first frame after the last post-water touch at which the hand has travelled more than half of its reach extent back from the spout, within 0.5 s of the last touch. **Windows:** For every trial the RESPONSE window was the 0.5 s following that outcome’s anchor and the REFERENCE window the 0.5 s immediately preceding it.

## Statistics

Non-parametric tests were used throughout: Mann–Whitney U for unpaired two-group comparisons, Wilcoxon signed-rank for paired comparisons, Fisher’s exact test for contingency tables, Spearman correlation for monotonic association, and χ² for larger contingency tables. Permutation and surrogate nulls are described with the analyses that use them; in every case the observed statistic was passed through the identical estimator as the surrogates. One-sided tests were used only where the hypothesis was directional, and this is stated in the text. Where an analysis compares populations of very different size, a matched-n control is reported alongside the full-sample test. All random procedures were seeded deterministically so that re-running the pipeline reproduces the reported values exactly.

Tests used for the transcriptomic analyses (DESeq2 Wald tests with Benjamini–Hochberg adjustment, GSEA, MetaNeighbor, binomial and Fisher’s exact tests) are described with the corresponding analyses.

## Software and data handling

Analyses were implemented in Python using NumPy, SciPy, pandas, scikit-learn and Matplotlib, with Suite2p for image processing, DeepLabCut 3.0.0 (PyTorch) for pose estimation, and CASCADE for spike inference. The analysis pipeline is orchestrated as an ordered set of stages sharing a single output base, with each stage reading the persisted outputs of its predecessors, so that cell selection and trial indexing are identical across analyses. Single-nucleus analyses used Cell Ranger 7.0.0, SoupX, scDblFinder, Seurat, Harmony, MapMyCells, DESeq2, clusterProfiler, limma and MetaNeighbor, gene-family, concordance and NDD figures were drawn in Python with Matplotlib.

## AI Use Declaration

Generative AI (Claude, Opus 5.5, Anthropic) was used to assist with the analysis code. The authors reviewed and verified all AI-assisted content and take full responsibility for the accuracy and integrity of the work.

## Author Contributions

JH and SMS conceived of the study. SMS, SS and JH designed experiments. SMS, JZ and SS performed the experiments. SMS and YZ analyzed data. HM helped with setting up of 2P behavioral rig. YL provided inputs on the RGD behavior. SZ made the AAVs. SMS, YZ and JH wrote the manuscript.

## Acknowledgements

We thank Dr. Xu An for helpful discussions. We thank Bao-Xia for her assistance with genotyping. Funding: NIH grant U19MH114823-01 (ZJH). NIH Director’s Pioneer Award 1DP1MH129954-01 (ZJH). International Postdoc Grant, Swedish Research Council grant no: 2021-00238 (SMS).

---

**Extended Data Fig. 1.**
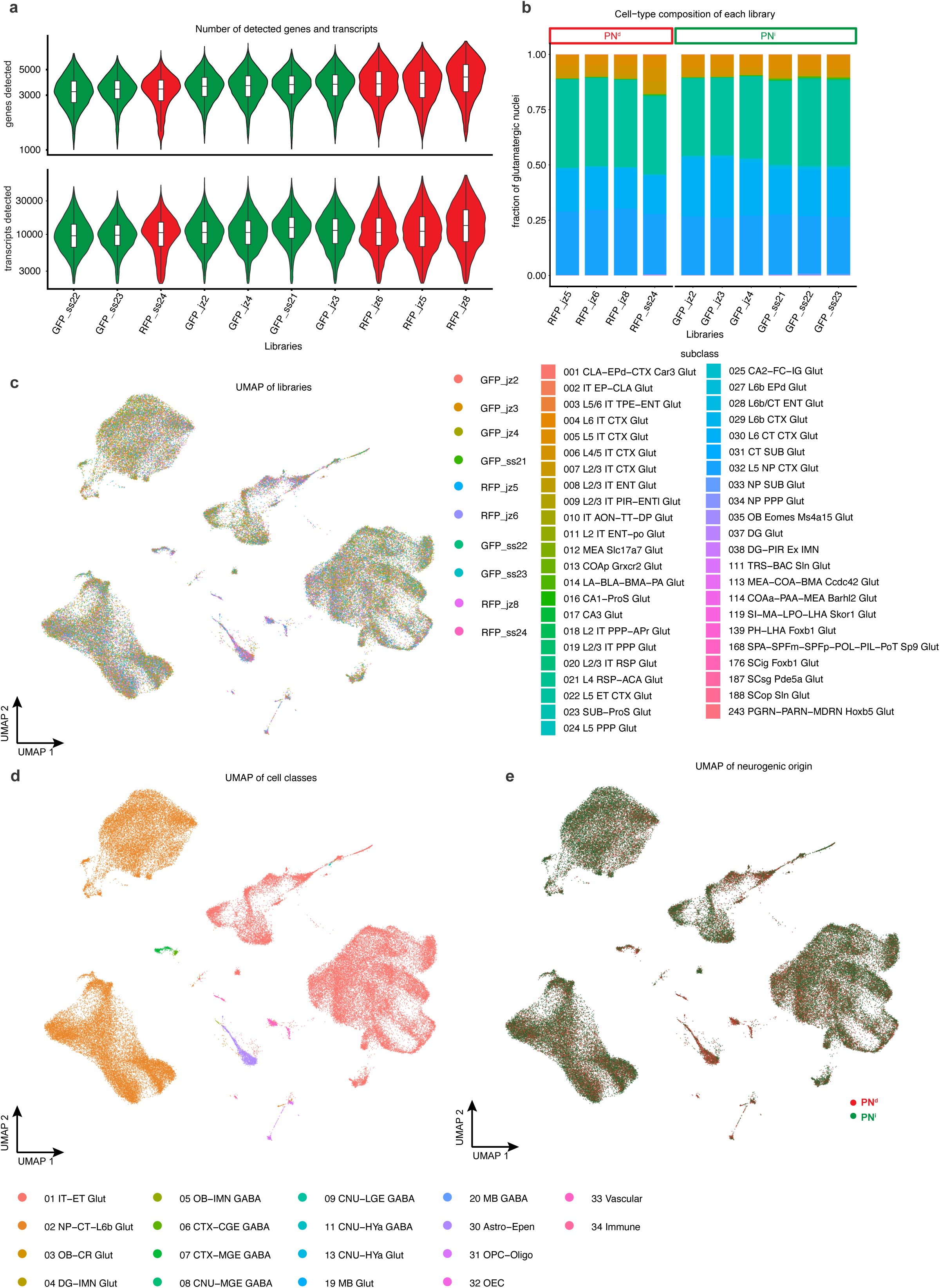
Quality and composition of the single-nucleus libraries. **a.** Number of detected genes and transcripts of each library, GFP (ET^i^) and RFP (ET^d^): genes and transcripts (UMIs) per nucleus. **b.** Cell-type composition of each library, grouped by dNG/iNG, colored by glutamatergic subclass. **c.** UMAP of all sequenced nuclei colored by library of origin. **d.** The same UMAP colored by cell class. **e.** The same UMAP colored by neurogenic origin (ET^d^ red, ET^i^ green); ET^d^ and ET^i^ nuclei intermingle throughout.

**Extended Data Fig. 2.**
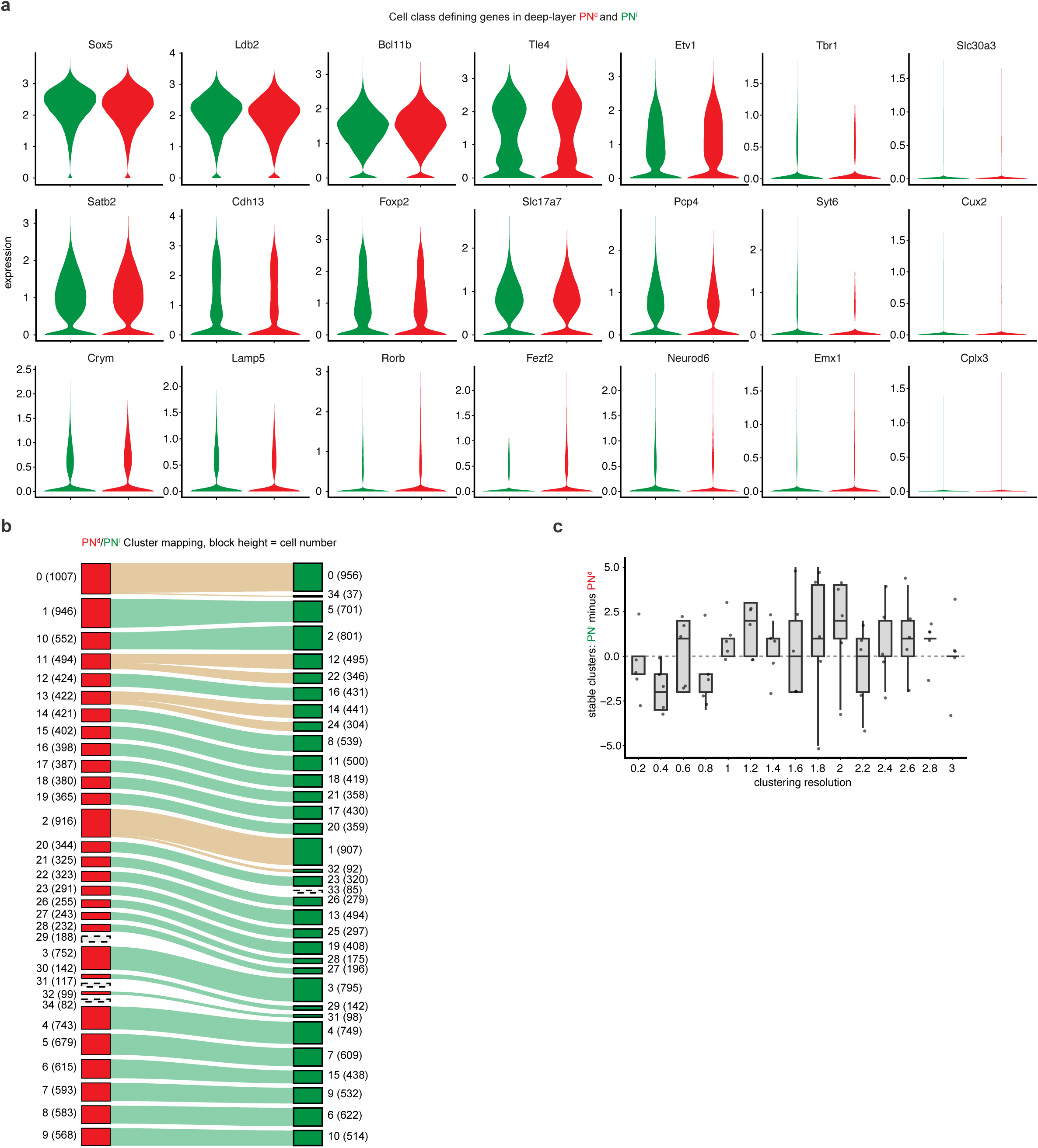
PN^d^ and PN^i^ express subclass-defining genes similarly and map onto one another across transcriptomic types. **a.** Expression of cell-class-defining genes in PN^d^ and PN^i^ nuclei. **b.** Mapping between independently clustered PN^d^ (left) and PN^i^ (right) nuclei, linked by expression similarity at resolution 2.8, the resolution with the largest total number of stable clusters across both lineages.; block height, number of nuclei (in parentheses), scaled within each neurogenic population. Green, one-to-one correspondence; tan, one ET^d^ cluster corresponding to several ET^i^ clusters; dashed, clusters without a counterpart. **c.** Difference in number of stable clusters (ET^i^ minus ET^d^) against clustering resolution.

**Extended Data Fig. 3.**
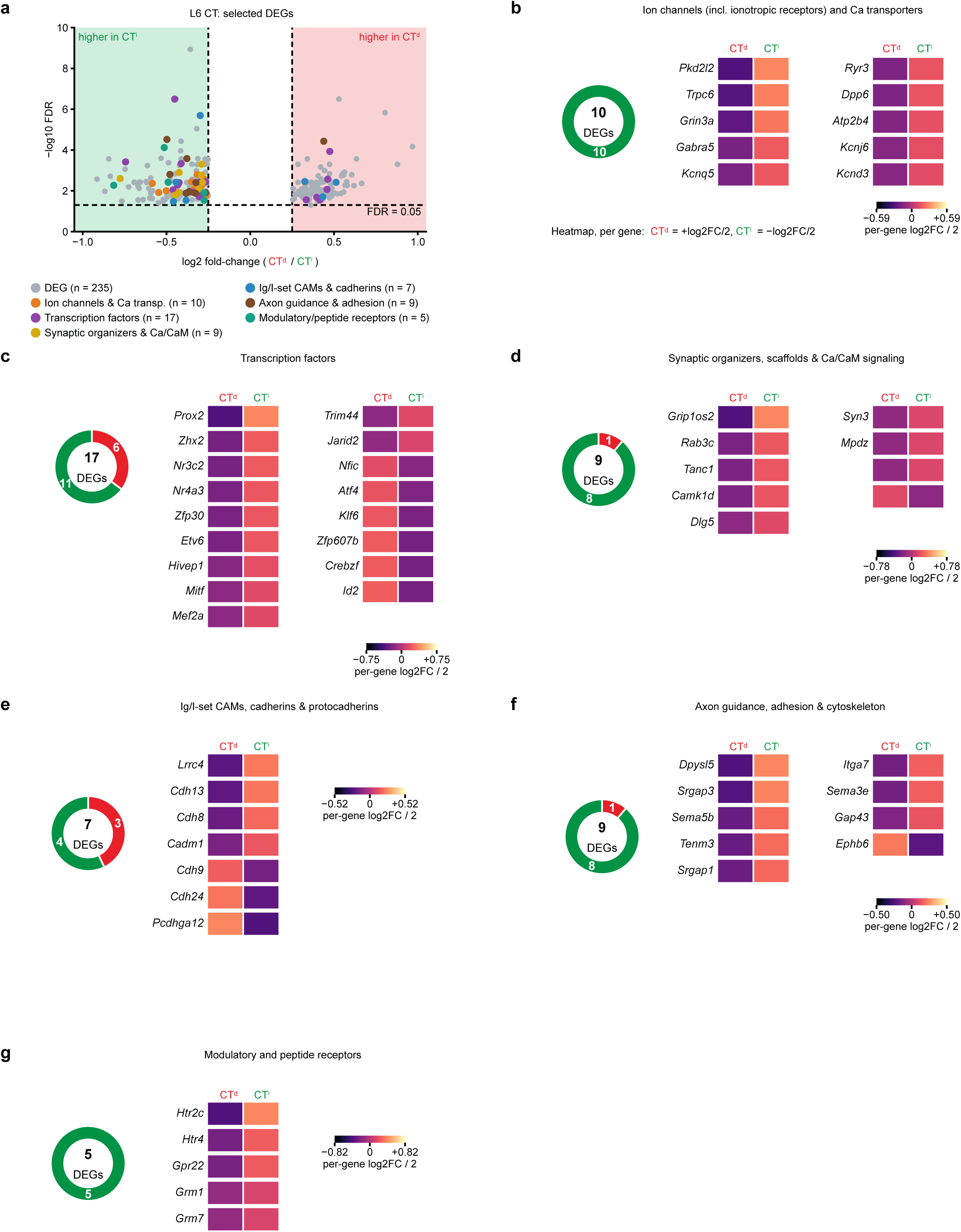
Single-gene differential expression across effector-gene families in L6 CT. **a.** Volcano of L6 CT DEGs (CT^d^/CT^i^; −log10 FDR versus log2 fold-change). Grey, all 235 DEGs; coloured, DEGs in the selected gene families (ion channels & Ca transporters n = 10, transcription factors n = 17, synaptic organizers & Ca/CaM n = 9, Ig/I-set CAMs & cadherins n = 7, axon guidance & adhesion n = 9, modulatory/peptide receptors n = 5). **b.** Ion channels (including ionotropic receptors) and Ca²⁺ transporters (10 DEGs, all higher in CT^i^). Donut of the direction split; per-gene heatmap (CT^d^ = +log2FC/2, CT^i^ = −log2FC/2). **c.** Transcription factors (17 DEGs; 11 higher in CT^i^, 6 in CT^d^); conventions as in b. **d.** Synaptic organizers, scaffolds and Ca²⁺/CaM signaling (9 DEGs; 8 higher in CT^i^, 1 in CT^d^); conventions as in b. **e.** Ig/I-set cell-adhesion molecules, cadherins and protocadherins (7 DEGs; 4 higher in CT^i^, 3 in CT^d^); conventions as in b. **f.** Axon guidance, adhesion and cytoskeleton (9 DEGs; 8 higher in CT^i^, 1 in CT^d^); conventions as in b. **g.** Modulatory and peptide receptors (5 DEGs, all higher in CT^i^); conventions as in b. Direction is the neurogenic population with higher expression. Gene families as in Fig. 2.

**Extended Data Fig. 4.**
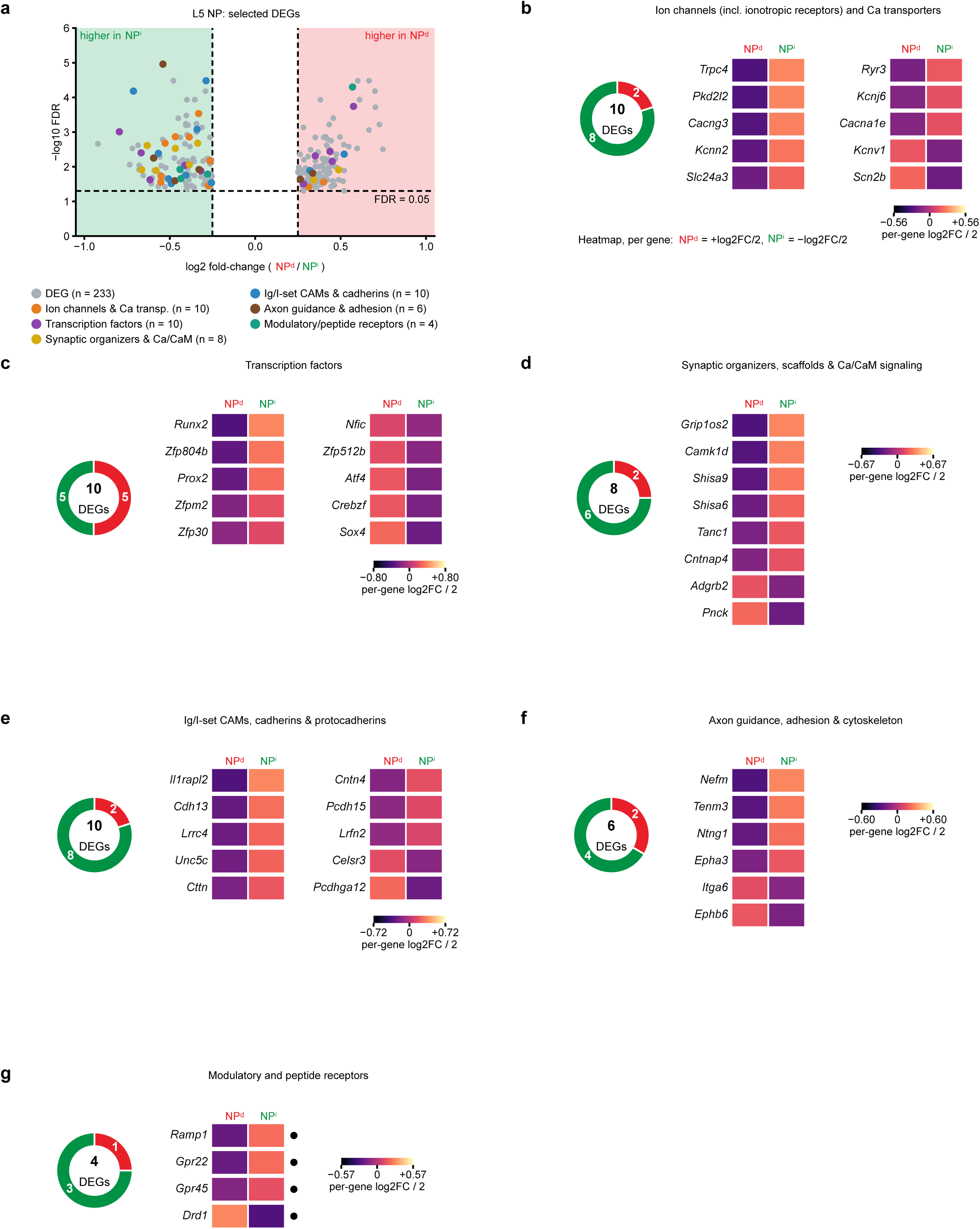
Single-gene differential expression across effector-gene families in L5 NP. **a.** Volcano of L5 NP DEGs (NP^d^/NP^i^; −log10 FDR versus log2 fold-change). Grey, all 233 DEGs; colored, DEGs in the selected gene families (ion channels & Ca transporters n = 10, transcription factors n = 10, synaptic organizers & Ca/CaM n = 8, Ig/I-set CAMs & cadherins n = 10, axon guidance & adhesion n = 6, modulatory/peptide receptors n = 4). **b.** Ion channels (including ionotropic receptors) and Ca²⁺ transporters (10 DEGs; 8 higher in NP^i^, 2 in NP^d^). Donut of the direction split; per-gene heatmap (NP^d^ = +log2FC/2, NP^i^ = −log2FC/2). **c.** Transcription factors (10 DEGs; 5 higher in NP^i^, 5 in NP^d^); conventions as in b. **d.** Synaptic organizers, scaffolds and Ca²⁺/CaM signaling (8 DEGs; 6 higher in NP^i^, 2 in NP^d^); conventions as in b. **e.** Ig/I-set cell-adhesion molecules, cadherins and protocadherins (10 DEGs; 8 higher in NP^i^, 2 in NP^d^); conventions as in b. **f.** Axon guidance, adhesion and cytoskeleton (6 DEGs; 4 higher in NP^i^, 2 in NP^d^); conventions as in b. **g.** Modulatory and peptide receptors (4 DEGs; 3 higher in NP^i^, 1 in NP^d^); conventions as in b. Direction is the neurogenic population with higher expression. Gene families as in Fig. 2.

**Extended Data Fig. 5.**
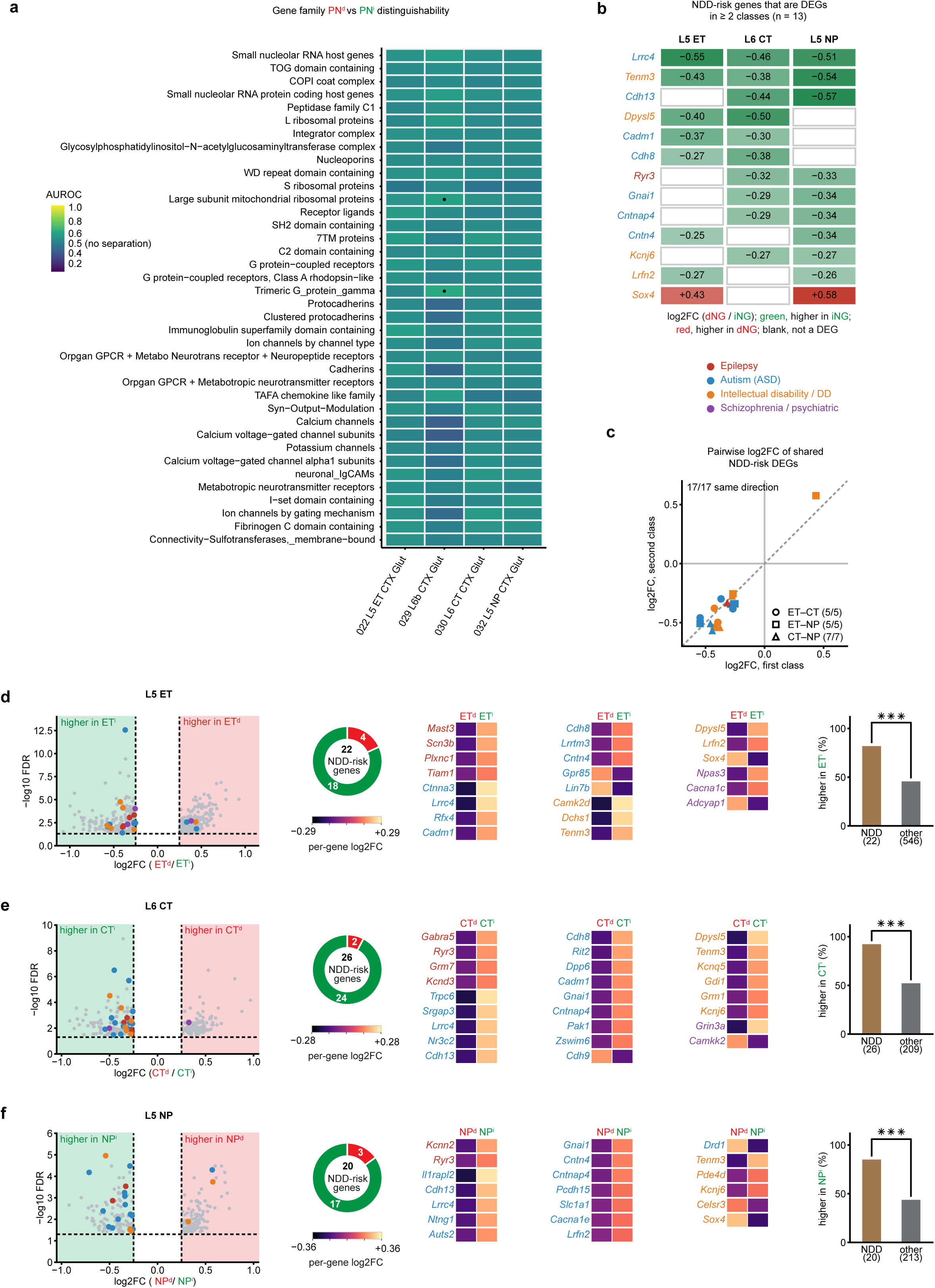
Gene families shift expression without classifying PN^d^ and PN^i^, and PN^i^-shifted genes are enriched for neurodevelopmental-disorder risk in all deep-layer PN subclasses. **a.** Classification of PN^d^ versus PN^i^ nuclei by each gene family (AUROC) in each deep-layer subclass; Dots, predictive families (FDR < 0.05, AUROC ≥ 0.6). Families in **Fig.1h** are shown here. AUROC close to 0.5 indicates no categorical separation despite significant family-level enrichment. **b.** NDD-risk genes that are DEGs in two or more classes (n = 13): log2FC (dNG/iNG) in L5 ET, L6 CT and L5 NP; green, higher in iNG; red, higher in dNG; blank, not a DEG in that class. Gene names colored by disorder (epilepsy, autism, intellectual disability/developmental delay, schizophrenia/psychiatric). **c.** Pairwise log2FC of the shared NDD-risk DEGs for each class pair (marker, pair; colour, disorder); all 17 pairwise comparisons are in the same direction (L5 ET–L6 CT 5/5, L5 ET–L5 NP 5/5, L6 CT–L5 NP 7/7). **d.** L5 ET. Left, volcano of DEGs with NDD-risk DEGs coloured by disorder (grey, other DEGs); middle, donut of the direction split (22 NDD-risk DEGs; 18 higher in ET^i^, 4 in ET^d^) and per-gene heatmaps (ET^d^ = +log2FC/2, ET^i^ = −log2FC/2), gene names coloured by disorder; right, percentage higher in ET^i^ for NDD-risk DEGs against the class’s other DEGs (82% versus 46%; two-sided Fisher exact test, odds ratio 5.4, p = 8.5 × 10^−4^). **e.** L6 CT, as d (26 NDD-risk DEGs; 24 higher in CT^i^, 2 in CT^d^; 92% versus 52%; odds ratio 11.0, p = 4.2 × 10^−5^). **f.** L5 NP, as d (20 NDD-risk DEGs; 17 higher in NP^i^, 3 in NP^d^; 85% versus 44%; odds ratio 7.3, p = 6.3 × 10^−4^). NDD-risk genes: neuronal genes with strong (SFARI 1 or S, Genes4Epilepsy, DDD, GenCC definitive or strong, or published Mendelian NDD) or limited (SFARI 2, GenCC moderate, single family or association) human genetic evidence; all are single-gene DEGs. ***, p < 0.001.

**Extended Data Fig. 6.**
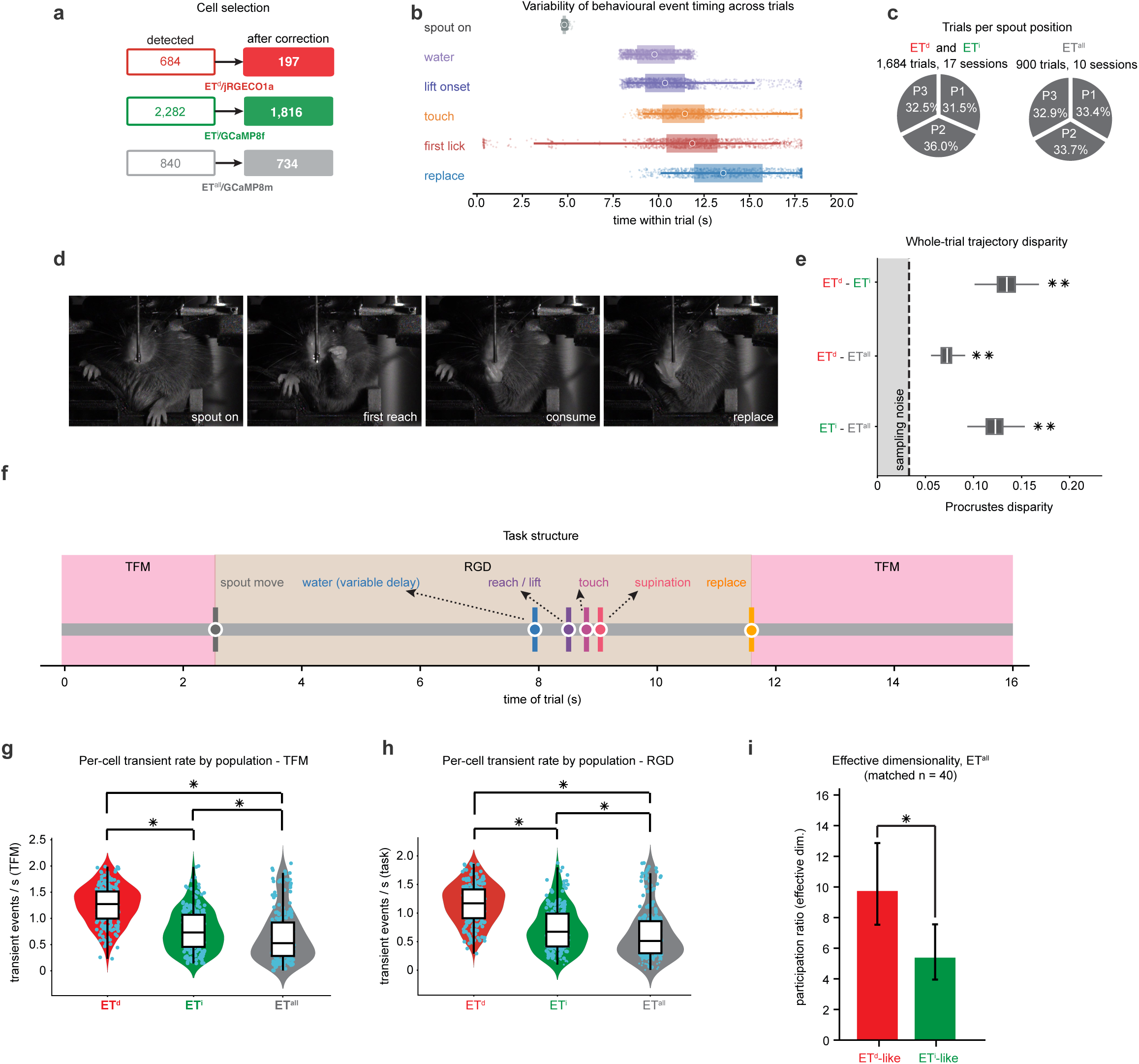
Cell selection, event timing, trajectory disparity, transient rates and ET^all^ dimensionality. **a.** Cell-selection pipeline: detected ROIs, dual-channel overlap removal giving 197 ET^d^ (*jRGECO1a*), 1,816 ET^i^ (*GCaMP8f*) and 734 ET^all^ (*GCaMP8m*) cells. **b.** Event-timing variability across trials during RGD (spout on, water, lift onset, touch, first lick and replace; boxes, IQR; points, trials). The motorised spout varied by 0.10 s (s.d.) against 2.10–3.47 s for animal-initiated events, over 1,684 trials. **c.** Trials per spout position (P1–P3) for the ET^d^/ET^i^ cohort (1,684 trials, 17 sessions) and the ET^all^ cohort (900 trials, 10 sessions). **d.** Example forelimb and face camera frames at spout on, first reach, consume and replace onset. **e.** Pairwise whole-trial trajectory disparity for ET^d^, ET^i^ and ET^all^ (Procrustes disparity over 200 draws at matched cell count, n = 197); ET^d^-ET^i^ 0.135, ET^d^-ET^all^ 0.073, ET^i^-ET^all^ 0.123; all clear their floor and null (permutation p = 0.005). ET^all^ lies between ET^d^ and ET^i^. **f.** Task structure schematic. Task-free movement (TFM) windows flank a reach-grasp-drink (RGD) window; markers indicate spout move, water (variable delay), reach/lift, touch, supination and replace. **g.** Per-cell TFM transient rate for ET^d^, ET^i^ and ET^all^. Populations subsampled to a common n = 197 over 50 draws; pairwise permutation test, Bonferroni-corrected, p = 0.015 for all three comparisons (the resolution limit of 200 permutations). Violins with box plots (median and IQR; whiskers 1.5 × IQR); points, cells; *, p < 0.015. **h.** As in **g** for the RGD (task) window (medians 1.17, 0.68 and 0.55 events/s). **i.** Effective dimensionality (participation ratio) of the whole-trial response manifold within ET^all^, for cells matched cell by cell to ET^d^ (ET^d^-like) or to ET^i^ (ET^i^-like; as in **Extended Data Fig. 7d,e**), at matched n = 40. Bars, participation ratio; error bars, 95% bootstrap intervals over matched-n draws; *, reference-shuffle permutation p = 0.015.

**Extended Data Fig. 7.**
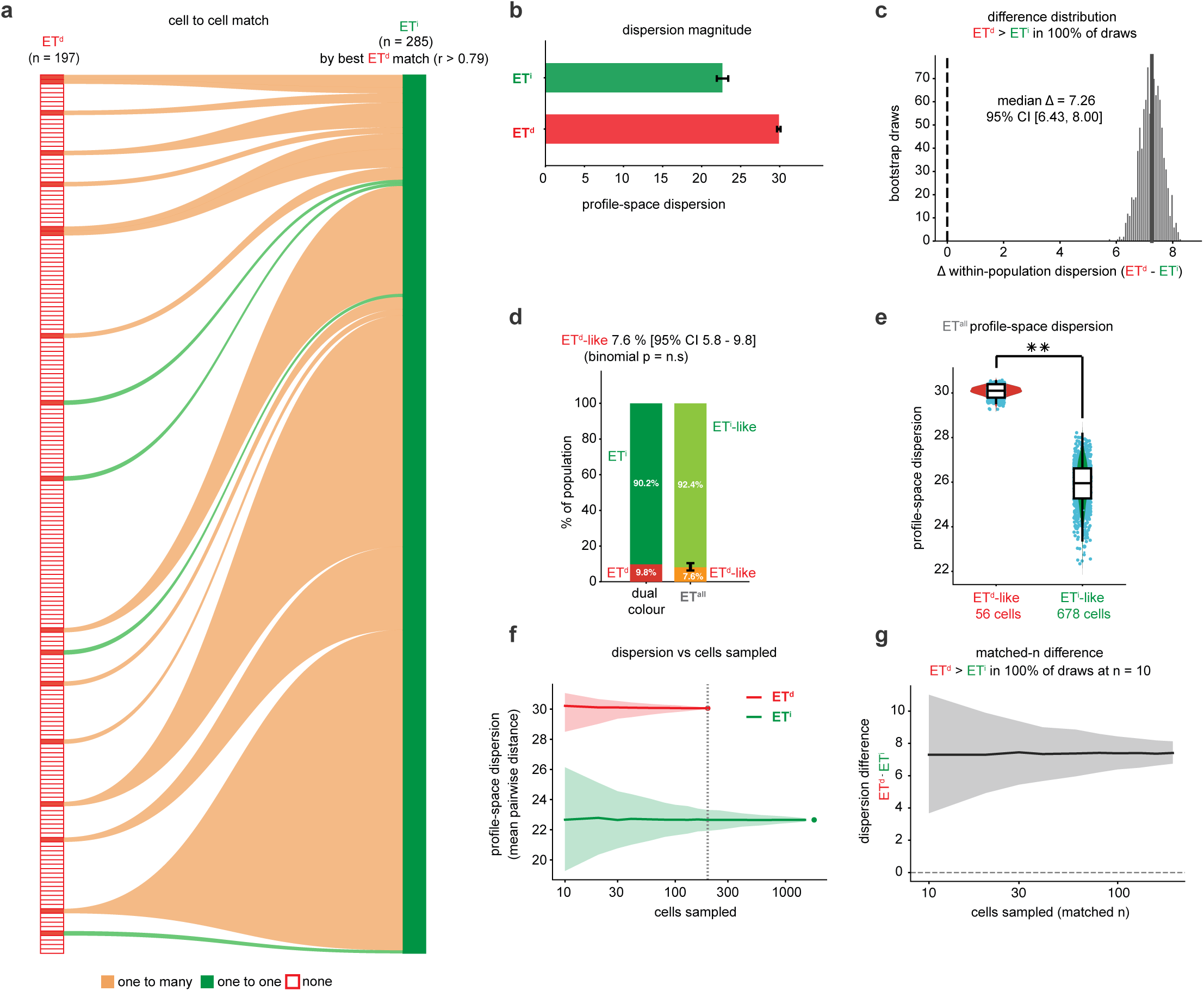
Cell-to-cell correspondence, dispersion and ET^all^ indicator controls. **a.** Cell-to-cell mapping over the RGD window (−2 to +10 s from spout onset). Each ET^i^ cell was assigned to the ET^d^ cell with the most similar trial-averaged profile (Pearson r); shown are the 285 of 1,816 ET^i^ cells with r > 0.79 (95th percentile of a circular time-shift null), which map onto 18 of 197 ET^d^ cells. Orange, ET^d^ cells receiving several ET^i^ cells (one to many; 14); green, ET^d^ cells receiving one (one to one; 4); open boxes, ET^d^ cells receiving none (179). **b.** Whole-population dispersion (mean pairwise Euclidean distance between z-scored whole-trial profiles): ET^d^ 29.9 versus ET^i^ 22.7 (matched-n bootstrap; error bars, 95% interval). **c.** Bootstrap distribution of the dispersion difference (ET^d^-ET^i^), median Δ = 7.26 (95% CI 6.43-8.00), positive in 100% of draws. **d.** ET^all^ composition from cell-to-cell matching: 56 of 734 cells (7.6%; 95% CI 5.8–9.8%) are ET^d^-like, against 9.8% ET^d^ measured by neurogenic labelling in the dual-color cohort (binomial p = 0.054). **e.** Profile-space dispersion of ET^all^ cells assigned cell by cell to a ET^d^ or ET^i^, ET^d^-matched (56 cells) against ET^i^-matched (678 cells), at matched n = 40 per draw; ET^d^-matched greater in 100% of draws (reference-shuffle permutation p = 0.005). **f.** Dispersion against the number of cells sampled for ET^d^ (n = 197) and ET^i^ (n = 1,816); 1,000 random subsets per size; line, median; shading, 95% interval; dots, full populations; dotted line, n = 197. **g.** Difference in dispersion (ET^d^-ET^i^) at matched subset size; line, median; shading, 95% interval. ET^d^ exceeds ET^i^ in 100% of draws at every size, including n = 10.

**Extended Data Fig. 8.**
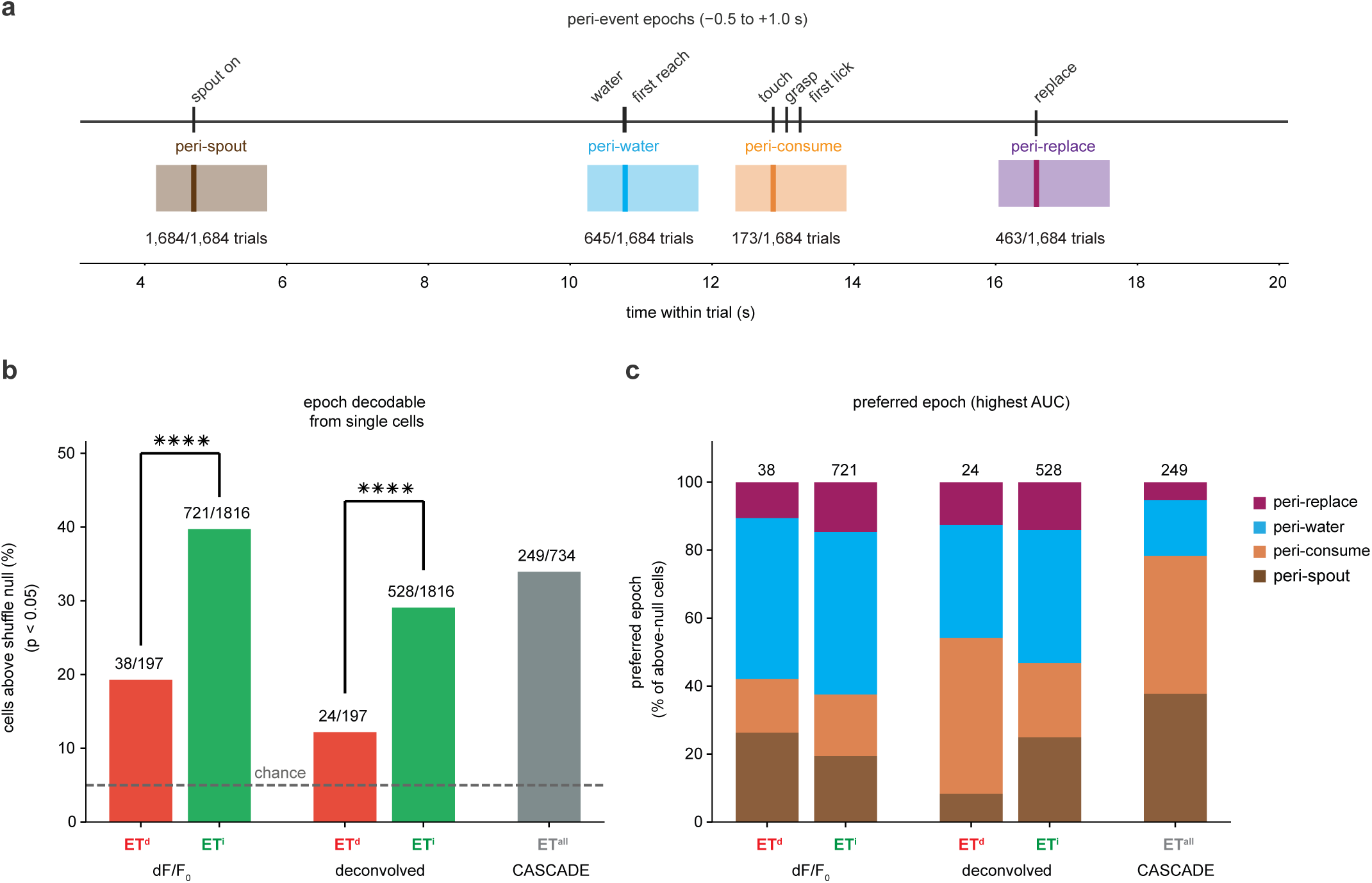
Single-cell peri-epoch decoding in ET^d^, ET^i^ and ET^all^. **a.** The four peri-event epochs (peri-spout, peri-water, peri-consume and peri-replace), each −0.5 to +1.0 s about its anchor (spout on, first reach after water delivery, first touch after water delivery and replace), with usable-trial counts (1,684, 645, 173 and 463 of 1,684 trials). **b.** Percentage of cells from which the four peri-epochs are decoded above a within-trial label-shuffle null (p < 0.05; leave-one-trial-out nearest-centroid classifier, 1,000 shuffles): ET^d^ and ET^i^ on dF/F_0_ (38/197, 19.3%, versus 721/1,816, 39.7%; Fisher exact p = 6.8 × 10^−9^) and on deconvolved spikes (24/197, 12.2%, versus 528/1,816, 29.1%; p = 8.6 × 10^−8^), and ET^all^ on CASCADE-inferred spike rates (249/734, 33.9%). Dashed line, chance (5%); ****, p < 10^−4^. **c.** Preferred epoch (highest AUC) of the above-null cells in each group; numbers, above-null cells.

**Extended Data Fig. 9.**
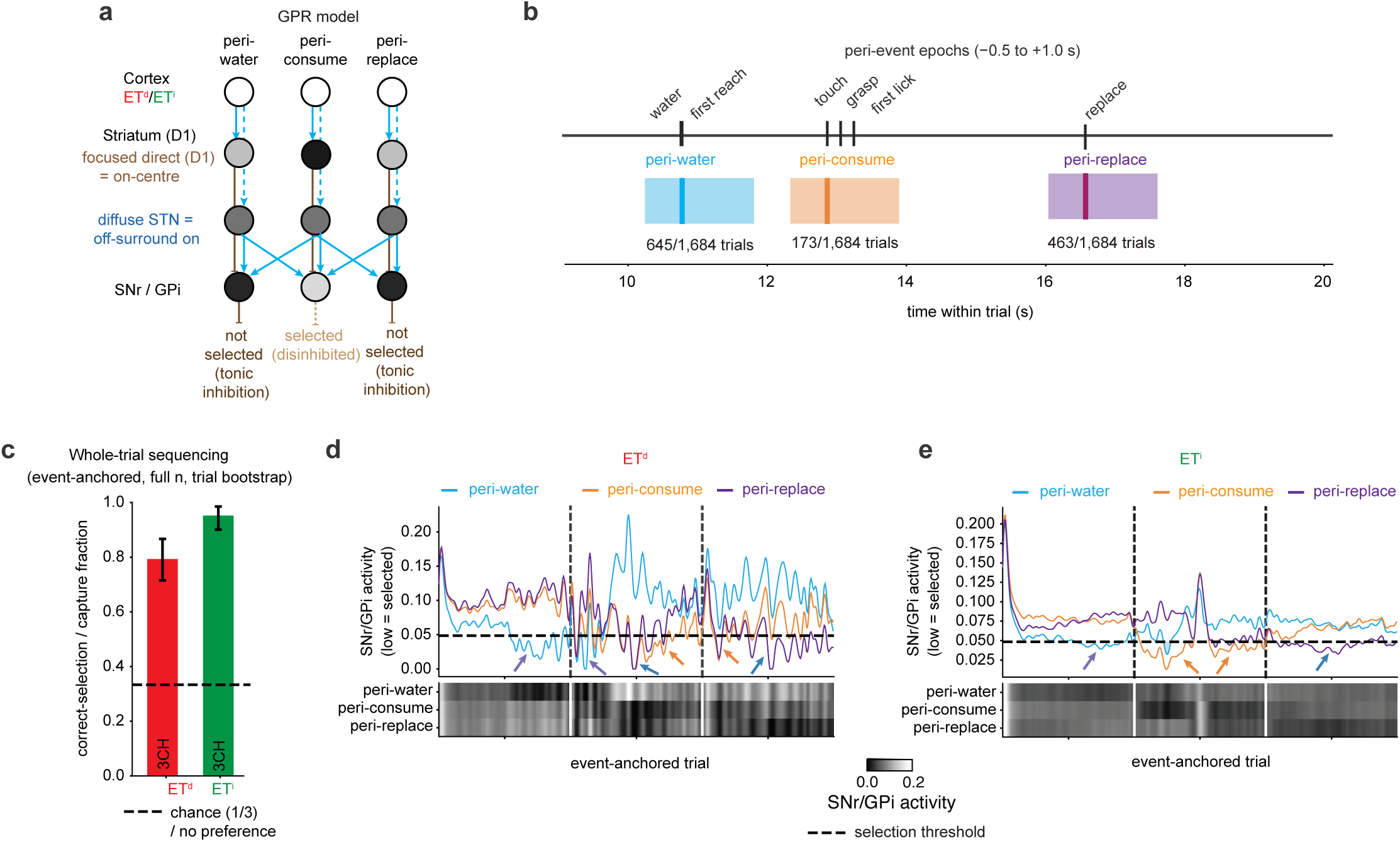
A basal-ganglia selection model temporally sequences RGD with ET^i^ activity. **a.** The GPR basal-ganglia model (see also^69^). Cortical ET^d^ or ET^i^ activity supplies the salience of three channels (peri-water, peri-consume, peri-replace). A focused direct (D1) pathway inhibits the output nucleus for its own channel (on-centre); the subthalamic projection excites it in proportion to summed salience (diffuse off-surround). **b.** Peri-event epochs supplying the three channels (peri-water, peri-consume and peri-replace), each −0.5 to +1.0 s about its anchor, with usable-trial counts. **c.** Whole-trial sequencing (event-anchored, full n, trial bootstrap): correct selection (3CH); dashed line, chance (1/3). ET^i^ 95.1% versus ET^d^ 79.3%. **d-e.** SNr/GPi activity (low = selected) when the whole **(d)** ET^d^ or **(e)** ET^i^ population drives the model; each channel’s salience is the population’s deconvolved activity, each cell weighted by its mean response in that peri-epoch. Below, SNr/GPi activity per channel across the event-anchored trial; dashed line, selection threshold. In ET^i^ each channel is selected in its own window in turn; in ET^d^ selection is fragmented.

**Extended Data Fig. 10.**
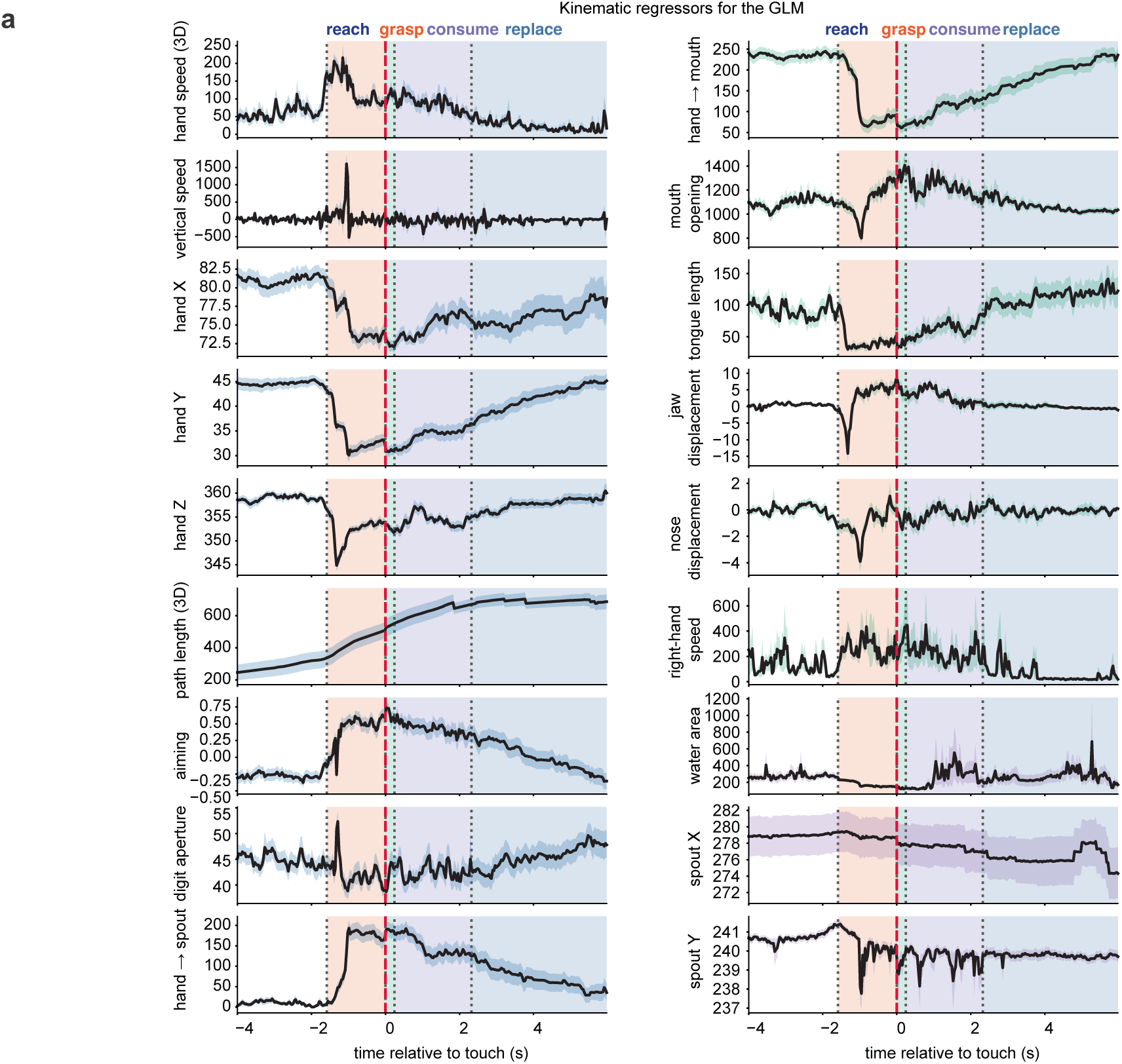
Kinematic regressors for the GLM. **a.** The 18 kinematic series entering the GLM, aligned to touch (mean ± s.e.m.), with reach, grasp, drink and replace epochs shaded: hand speed (3D), vertical speed, hand X/Y/Z, path length, aiming, digit aperture, hand → spout, hand → mouth, mouth opening, tongue length, jaw and nose displacement, right-hand speed, water area and spout X/Y.

**Extended Data Fig. 11.**
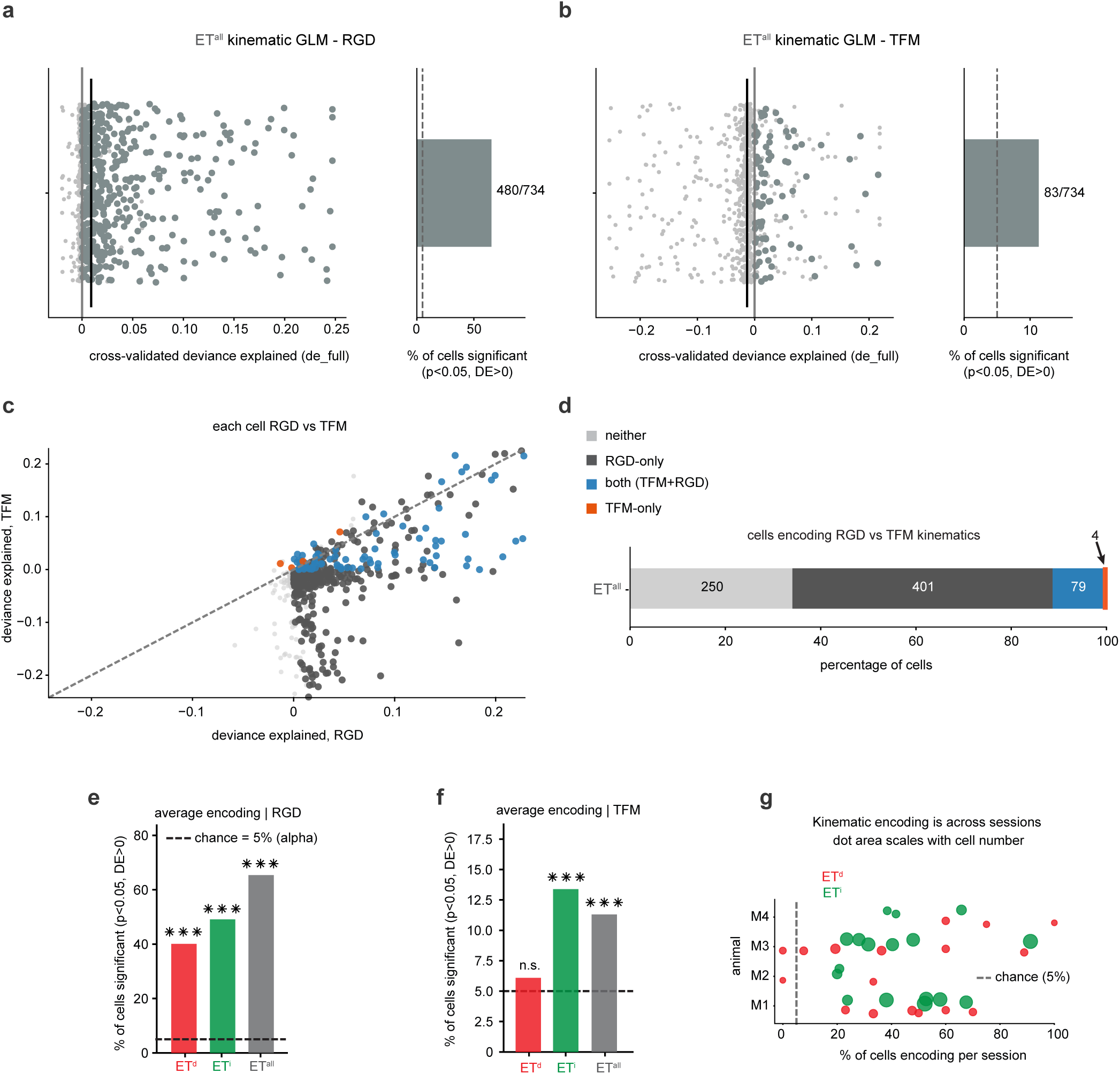
ET^all^ kinematic GLM and encoding controls. **a.** Per-cell kinematic GLM in ET^all^ (n = 734), RGD: deviance explained per cell (left) and percentage of cells encoding (right; 480/734, 65.4%); conventions as in Fig. 6a. **b.** As in **a** for TFM (83/734, 11.3%). **c.** Deviance explained per ET^all^ cell in RGD against TFM, colored by encoding category as in d; dashed line, identity. **d.** Encoding category per ET^all^ cell: neither 250, RGD only 401, both 79 and TFM only 4. **e.** Percentage of cells encoding in RGD for ET^d^, ET^i^ and ET^all^; dashed line, chance (5%); ***, binomial p < 0.001 against chance. **f.** As in **e** for TFM; ET^d^ at chance (n.s.), ET^i^ and ET^all^ above. **g.** Per-session percentage of cells encoding, ET^d^ and ET^i^, grouped by animal; dot area scales with cell number; dashed line, chance (5%).

**Extended Data Fig. 12.**
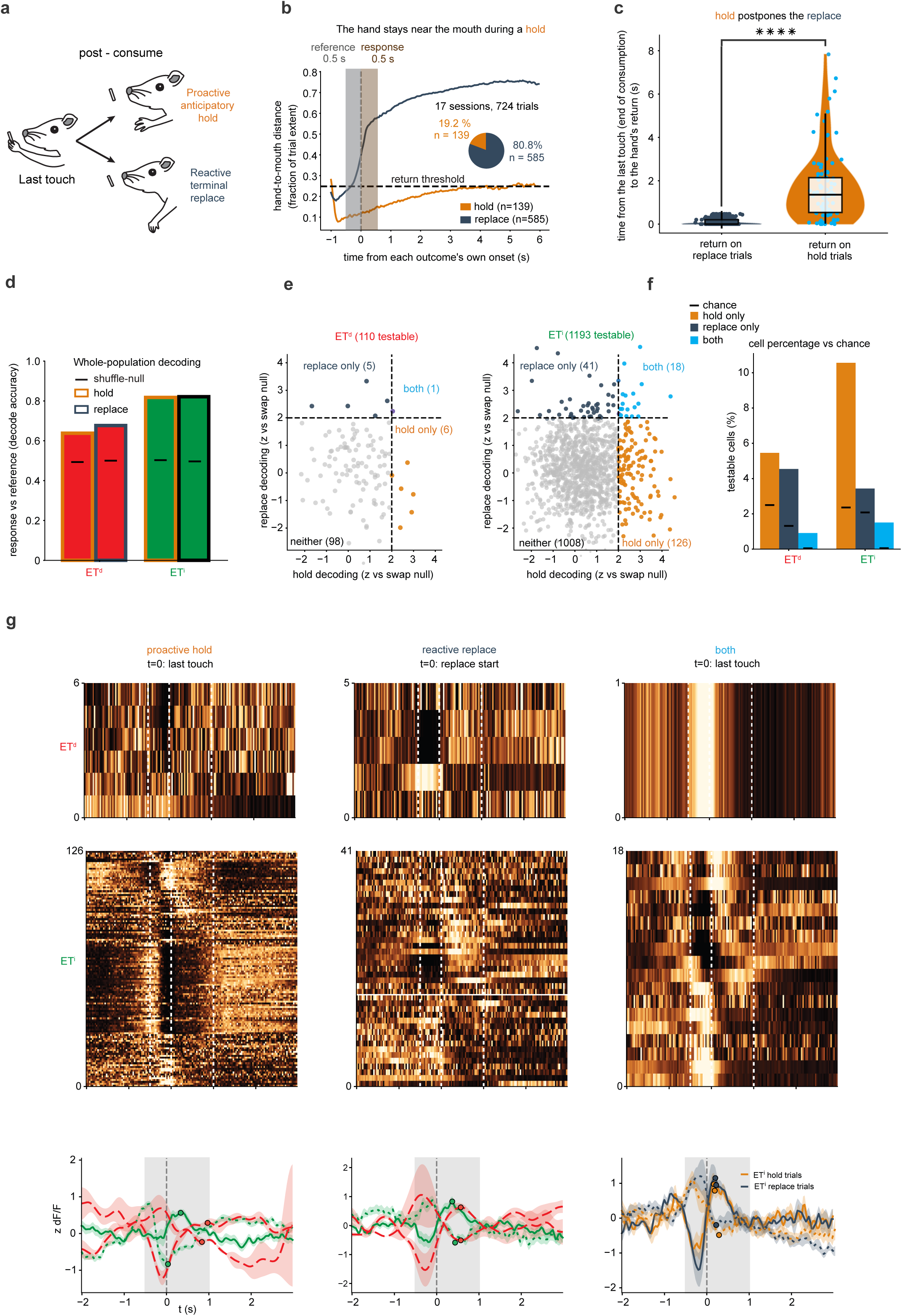
ET^i^ forms separate output channels for a proactive anticipatory and a reactive terminal ending of the RGD action. **a.** After consuming, the animal either keeps the hand raised (proactive anticipatory hold) or returns it to the perch (reactive terminal replace). **b.** Hand-to-mouth distance about each ending’s own onset, normalized within trial; dashed line, return threshold. Shaded bands mark the two 0.5 s windows used for the decoding analysis (panels **d-g**): the reference window, the 0.5 s immediately before onset, and the response window, the 0.5 s immediately after; the two are identical in width and placement for both endings and differ only in the anchoring event. Inset, frequency of the two endings among 724 analyzed trials (139 hold, 19.2%; 585 replace, 80.8%). **c.** Time from the last touch until the hand returns, on replace and hold trials; a hold significantly postpones the replace (Mann–Whitney, one-sided; ****, p < 0.0001).rr **d.** Whole-population decoding of each ending against its reference window for ET^d^ and ET^i^ (ET^d^ 0.638 hold, 0.677 replace; ET^i^ 0.817 and 0.821); bars, decoding accuracy; horizontal lines, shuffle nulls (about 0.50). **e.** Single-cell hold and replace decoding, each as z against the cell’s own within-trial swap null, for testable cells (at least five trials of each ending; ET^d^ 110, ET^i^ 1,193); dashed lines, z = 2. ET^d^: hold only 6, replace only 5, both 1, neither 98; ET^i^: hold only 126, replace only 41, both 18, neither 1,008. **f.** Percentage of testable cells in each class against chance (black lines, rate in swapped-label draws): ET^i^ hold only 10.6% (chance 2.4%), replace only 3.4% (2.1%), both 1.5% (0.1%); ET^d^ hold only 5.5% (2.5%), replace only 4.5% (1.3%), both 0.9%. **g.** Trial-averaged activity of every cell in each class: hold only (t = 0, last touch; hold trials), replace only (t = 0, replace start; replace trials) and both (t = 0, last touch; hold and replace trials), for ET^d^ (top) and ET^i^ (middle); rows z-scored, cells whose activity rises at the ending above those whose activity falls (direction from the decoder windows, 0 to +0.5 s against −0.5 to 0 s). ET^i^ hold only, 47 rising and 79 falling; replace only, 25 and 16; both, 8 and 10. Bottom, mean ± s.e.m. of rising (solid) and falling (dotted) cells: hold only and replace only, ET^i^ green and ET^d^ red (dashed); both, ET^i^ on hold trials (orange) and replace trials (blue). Grey band, −0.5 to +1.0 s. Statistics. Non-parametric tests throughout: Mann–Whitney U (unpaired), Wilcoxon signed-rank (paired), Fisher’s exact test (contingency), Spearman correlation (association) and χ² (larger contingency tables); permutation and circular-shift nulls are specified per analysis; matched-n bootstrap is used for size-mismatched comparisons; one-sided tests are indicated where used. DEGs are considered significant if they have FDR < 0.05, |log2FC| ≥ 0.25, and their direction is consistent across batches and sex-balanced draws. Family-level enrichment is by gene-set enrichment analysis; consistently shifted genes (CSGs) are read by direction and consistency; gene-family panels show, for L5 ET (Fig. 2c**-h**), the CSGs that are also single-gene DEGs and, for L6 CT and L5 NP (**Extended Data Figs. 3 and 4**), the DEGs of each family. Unless stated otherwise, n = 197 ET^d^, 1,816 ET^i^ and 734 ET^all^ cells, from 17 + 10 sessions in 4 + 3 animals and 1,684 trials.

